# Structural Organization of a Type III-A CRISPR Effector Subcomplex Determined by X-ray Crystallography and Cryo-EM

**DOI:** 10.1101/452599

**Authors:** Bryan W. Dorsey, Lei Huang, Alfonso Mondragón

## Abstract

Clustered regularly interspaced short palindromic repeats (CRISPR) and their associated Cas proteins provide an immune-like response in many prokaryotes against extraneous nucleic acids. CRISPR-Cas systems are classified into different classes and types. Class 1 CRISPR-Cas systems form multi-protein effector complexes that includes a guide RNA (crRNA) used to identify the target for destruction. Here we present crystal structures of *Staphylococcus epidermidis* Type III-A CRISPR subunits Csm2 and Csm3 and a 5.2 Å resolution single-particle cryo-electron microscopy (cryo-EM) reconstruction of an effector subcomplex including the crRNA. The structures help to clarify the quaternary architecture of Type III-A effector complexes, as well as to provide details on crRNA binding, target RNA binding and cleavage, and intermolecular interactions essential for effector complex assembly. The structures allow a better understanding of the organization of Type III-A CRISPR effector complexes as well as highlighting the overall similarities and differences with other Class 1 effector complexes.

## Introduction

Many prokaryotes, including most archaea and many bacteria, utilize immune-like response systems to defend themselves against phages and genomic invasions. One of these adaptive systems is comprised of genomic loci formed by <u>c</u>lustered <u>r</u>egularly <u>i</u>nterspaced <u>s</u>hort <u>p</u>alindromic <u>r</u>epeats (CRISPRs) and their CRISPR associated (Cas) proteins (Horvath & Barrangou, 2010; Karginov & Hannon, 2010; Marraffini, 2015; Marraffini & Sontheimer, 2010; van der Oost, Westra, Jackson, & Wiedenheft, 2014b). Although originally CRISPR was thought to function as an antisense system targeting invading phages (Barrangou et al., 2007), it was later shown that it can also protect against horizontal gene transfer (Marraffini & Sontheimer, 2008) and transformation (van der Oost et al., 2014b). CRISPR loci contain several non-adjacent direct DNA repeats separated by variable sequences, called spacers, which correspond to segments of captured viral and plasmid sequences (Jansen, van Embden, Gaastra, & Schouls, 2002). These loci allow for immune memory against invading foreign genetic elements by integrating the foreign DNA, which is then transcribed and processed to form CRISPR RNA (crRNA) (Barrangou et al., 2007; Bolotin, Quinquis, Sorokin, & Ehrlich, 2005; Pourcel, Salvignol, & Vergnaud, 2005). The crRNA, in conjunction with Cas proteins, are utilized for the identification and eventual destruction of recurrent invaders (Koonin, Makarova, & Zhang, 2017).

There is a wide diversity in the organization and architecture of CRISPR-Cas systems. Two main classes (Class 1 and 2) have been identified from extensive comparative analysis of both the loci and proteins associated with each system (Makarova et al., 2015). Class 1 CRISPR-Cas systems (Types I, III, and IV) consist of multi-protein effector complexes that function together for target surveillance and defense, whereas Class 2 CRISPR-Cas systems (types II, V, and VI) have a single protein that incorporates both surveillance and defense stages into one protein, Cas9 or homologues (Koonin et al., 2017; Makarova et al., 2015). Within Class I systems, the number of proteins involved and the general architecture of the effector complexes varies and even within each type there are differences in subunit stoichiometry. Type I systems can be subdivided into seven subtypes whereas Type III CRISPR systems are further divided into four subtypes, III-A to III-D (Makarova et al., 2015). All Type III subtypes include the Type III-defining *cas10* gene (Makarova et al., 2011), which encodes the signature Type III Cas10 protein that is responsible for DNA target degradation (Ramia, Spilman, et al., 2014). There are two distinct effector complexes formed within Type III subtypes, known as the Csm (Types III-A/D) and Cmr (Types III-B/C) effector complexes (Rouillon et al., 2013; Zhang, Graham, Tello, Liu, & White, 2016; Zhang et al., 2012). While originally it was not clear whether Type III systems targeted foreign DNA or RNA for destruction, it was later shown that in Type III-A systems, both RNA and DNA can be cleaved, but utilize different active sites and in a co-transcriptional manner (Samai et al., 2015). Further studies showed that both Type III-A and III-B effector complexes act as RNases and target RNA-activated DNA nucleases (Elmore et al., 2016; Estrella, Kuo, & Bailey, 2016; Kazlauskiene, Tamulaitis, Kostiuk, Venclovas, & Siksnys, 2016), unifying the previously incongruent mechanisms of the Type III systems into one cohesive mechanism.

The S*taphylococcus epidermidis* RP62a CRISPR-Cas system has served as the paradigm for the Type III-A subtype. Only one CRISPR loci is encoded in *S. epidermidis* RP62a and contains three 33-34 base pair (bp) spacers flanked by four 37 bp direct repeats (**Figure 1A**) (Grissa, Vergnaud, & Pourcel, 2007). Previous studies have determined that the CRIPSR-Cas effector complex is composed of the Cas proteins Cas10, Csm2, Csm3, Csm4, and Csm5 in an undetermined stoichiometry and known as the Cas10-Csm complex (**Figure 1A**) (Hatoum-Aslan, Samai, Maniv, Jiang, & Marraffini, 2013). These proteins combine to form a two helical protein arrangement, known as the major and minor filaments (**Figure 1B**) (R. H. J. Staals et al., 2013; Tamulaitis, Venclovas, & Siksnys, 2017). The major filament is composed of multiple copies of Csm3, a RAMP (repeat-associated mysterious proteins) protein containing an RNA recognition motif (RRM) fold. The filament is capped on each end by other RAMPs, Csm4 and Csm5 (R. H. Staals et al., 2014; R. H. J. Staals et al., 2013; van der Oost, Westra, Jackson, & Wiedenheft, 2014a). Csm3, in particular, has been implicated in binding the crRNA guide, as well as functioning as an endoribonuclease that acts to cleave the ssRNA target in a periodic manner (Hatoum-Aslan et al., 2013). The minor filament is composed of multiple copies of Csm2, the smallest subunit in the Cas10-Csm complex. Its central function is thought to be in target strand binding (Tamulaitis et al., 2017). The protein component stoichiometry of both the major and minor filaments is dependent on the length of the crRNA, suggesting that the crRNA can modulate the assembly of the effector complex (Spilman et al., 2013; R. H. J. Staals et al., 2013). The final and largest component of the Cas10-Csm effector complex is the signature Type III protein Cas10, an exonuclease implicated in DNA target strand degradation (Jung et al., 2015; Ramia, Tang, Cocozaki, & Li, 2014). Thus, the Cas10-Csm complex contains two different catalytic activities, with DNA degradation achieved by Cas10 and RNA degradation accomplished by Csm3, with both guided by the crRNA.

**Figure 1.**
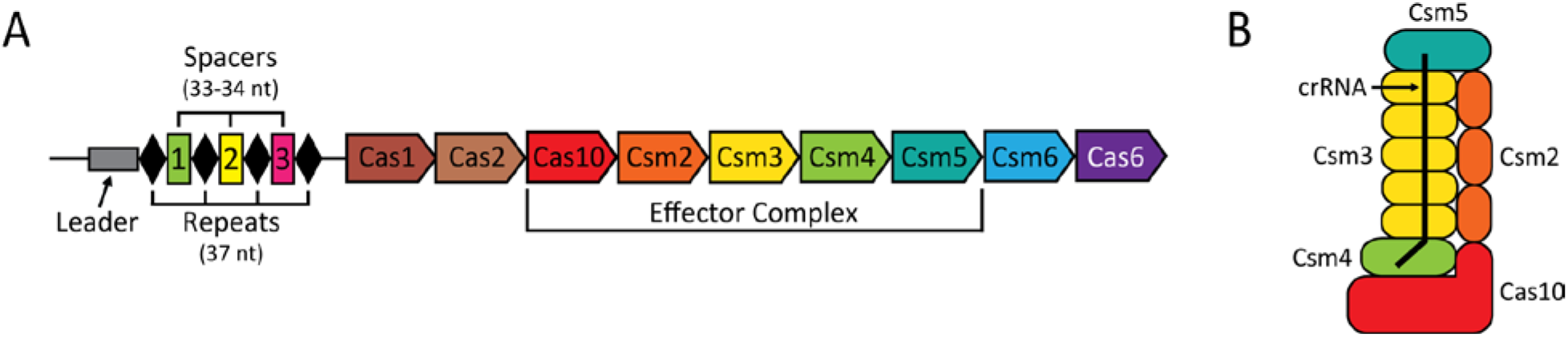
*S. epidermidis* CRISPR-Cas Locus and Effector Complex Schematics. (**A**) Schematic of *S. epidermidis* RP62a CRISPR-Cas locus architecture. The locus is composed of the leader (gray rectangle) and tandem copies of the repeats (black diamonds) and spacers (numbered 1-3), followed by the nine associated protein-coding *cas* genes. Five proteins (Cas10, Csm2-5) associate to form the effector complex. (**B**) Schematic diagram of a proposed model of the Type III-A CRISPR effector complex. The model is composed of five copies of Csm3 (yellow), three copies of Csm2 (orange), and one copy each of Csm4 (green), Csm5 (teal), and Cas10 (red), in addition to a guide CRISPR RNA (crRNA) of various lengths (black line) (Tamulaitis et al., 2017).

Our understanding of the architecture and mechanism of Type III-A systems has been limited due to the lack of high resolution structural information on the Cas10-Csm complex and its components. Although there is a wealth of information on other CRISPR types, such as Type I, the inherent diversity in the mechanism and architecture of the CRISPR systems precludes extrapolating from one type to the other, limiting how much we can learn from one type about another. To address this limitation, we have studied the structure of the *S. epidermidis* Cas10-Csm effector complex by a combination of crystallographic and cryo-electron microscopy (cryo-EM) studies. The studies reveal the crystal structures of Csm2 and Csm3, the two most abundant subunits forming the backbone of the complex. In addition, a 5.2 Å resolution cryo-EM structure of a subcomplex including most of the subunits and the crRNA sheds light on the overall architecture of the Cas10-Csm complex, the binding of crRNA in the complex, and the active site of Csm3. Furthermore, these structural studies help understand many of the previous biochemical observations on the mechanism of this particular CRISPR-Cas complex. Finally, there is a practical interest in understanding the mechanism of CRISPR-Cas immunity in *S. epidermidis*, as this gram positive bacteria are the most common of the coagulase-negative staphylococci found on the human epithelia (Otto, 2009, 2012) and is among the most common pathogens that lead to hospital-acquired nosocomial infections (Otto, 2012; von Eiff, Peters, & Heilmann, 2002).

## Results

### The crystal structure of *S. epidermidis* Csm2 reveals a structural organization intermediate between Type III-A and III-B molecules

*S. epidermidis* Csm2 (SeCsm2) was purified using a His-tag to apparent homogeneity under high salt conditions to help solubilize the purified protein **(Figure 2 - Figure supplement 1A**). Initial crystallization trials were performed with and without the affinity tag attached, with crystals of SeCsm2 only forming when the affinity tag was present (**Figure 2 - Figure supplement 1B**). Structural determination of SeCsm2 was accomplished by experimental Single Anomalous Dispersion (SAD) from selenomethionine-incorporated crystals (**Materials and Methods**). The final model of SeCsm2 (R_work_/R_free_ = 24.5%/29.4%) was obtained from data to 2.75 Å and consists of residues 14-28 and 37-139 with an additional 9 residues N-terminal to the start methionine identified as the tobacco etch virus (TEV) protease cleavage site (**Supplementary Table 1** and **Figure 2 - Figure supplement 1C**). This additional region provided essential crystal contacts in the crystal lattice (**Figure 2 - Figure supplement 1D**). After the structure was solved, the gene annotation for *csm2* was updated (Tatusova, Ciufo, Fedorov, O’Neill, & Tolstoy, 2015), showing an additional 13 residues upstream of the original starting residue. The new N-terminal region is predicted to be unstructured (**Figure 2 - Figure supplement 1E**), which could provide similar crystal contacts to the TEV protease cleavage site found in the crystal structure. It is presently unknown whether these additional N-terminal residues are relevant to the function of SeCsm2. All residue numbers stated in this work reflect the updated annotation for the *csm2* gene.

The overall structure of SeCsm2 reveals an exclusively α-helical fold composed of six helices (**Figure 2A**) forming a compact five-helix bundle with the sixth helix, α_4_, connecting two of the helices in the bundle. The organization of the helices generally follows the established fold for the small subunits of Class 1 CRISPR effector complexes (Gallo et al., 2016; Venclovas, 2016). There are two notable structural differences between the SeCsm2 structure and the only other Type III-A Csm2 crystal structure available, from *Thermotoga maritima* (TmCsm2; PDB: 5AN6) (Gallo et al., 2016) (**Figure 2B**). The first difference occurs in the loop between the first two helices, α_1_ and α_2_. In SeCsm2, there is an extended loop (Loop 1) consisting of 20 amino acids (residues 25-44) with the middle 8 amino acids (residues 29-36) disordered in the crystal (**Figure 2 - Figure supplement 1C**). This is much longer than the loop in TmCsm2, which only contains 6 amino acids (residues 30-35). In SeCsm2, Loop 1 contains many positively charged residues (K29, R31, K32, K34, and K36) and a pair of negatively charged residues (D30 and E35). Secondary structure predictions show an extension of α_1_ into Loop 1, however these residues are not part of α_1_ in the crystal (**Figure 2 - Figure supplement 1C**). This suggests that this region is more flexible than predicted and is consistent with the absence of electron density for some of this loop. However, it is possible that this highly positively charged loop could play a functional role during target strand binding. Additionally, the crystal packing reveals that Loop 1 in each molecule faces the solvent space (**Figure 2 - Figure supplement 1D**). The second difference involves the third helix in the bundle, α_3_, which rearranges in SeCsm2 to form two helices, α_4_ and α_5_, whereas it is a single helix in an unswapped monomer of TmCsm2 (**Figure 2B**) (Venclovas, 2016). The predicted secondary structure of SeCsm2 indicates that α_4_-α_5_ should form a single helix (**Figure 2 - Figure supplement 1C**), which would be consistent with the helix topology described for Type III-A structures. Instead, it breaks and kinks in the vicinity of residue 103. Interestingly, the occurrence of two helices at this position is more prevalent in the Type III-B small subunit, Cmr5 (Venclovas, 2016). Comparison of SeCsm2 α_4_-α_5_ to TmCsm2 α_3_ and α_4_-α_5_ from the Type III-B *Thermus thermophilus* HB8 Cmr5 structure (TtCmr5; PDB: 2ZOP) (Sakamoto et al., 2010) shows that SeCsm2 adopts an intermediate structural conformation between the single, linear helix in TmCsm2 and the pair of near perpendicular helices in TtCmr5 (**Figure 2C**). This ‘intermediate structure’ observation can be extended to the entire SeCsm2, with its relative helix orientations more similar to Type III-A small subunits, but including helix topologies reminiscent of those found in Type III-B small subunits. This suggests that the small subunits form a more diverse group of structures than previously thought.

**Figure 2.**
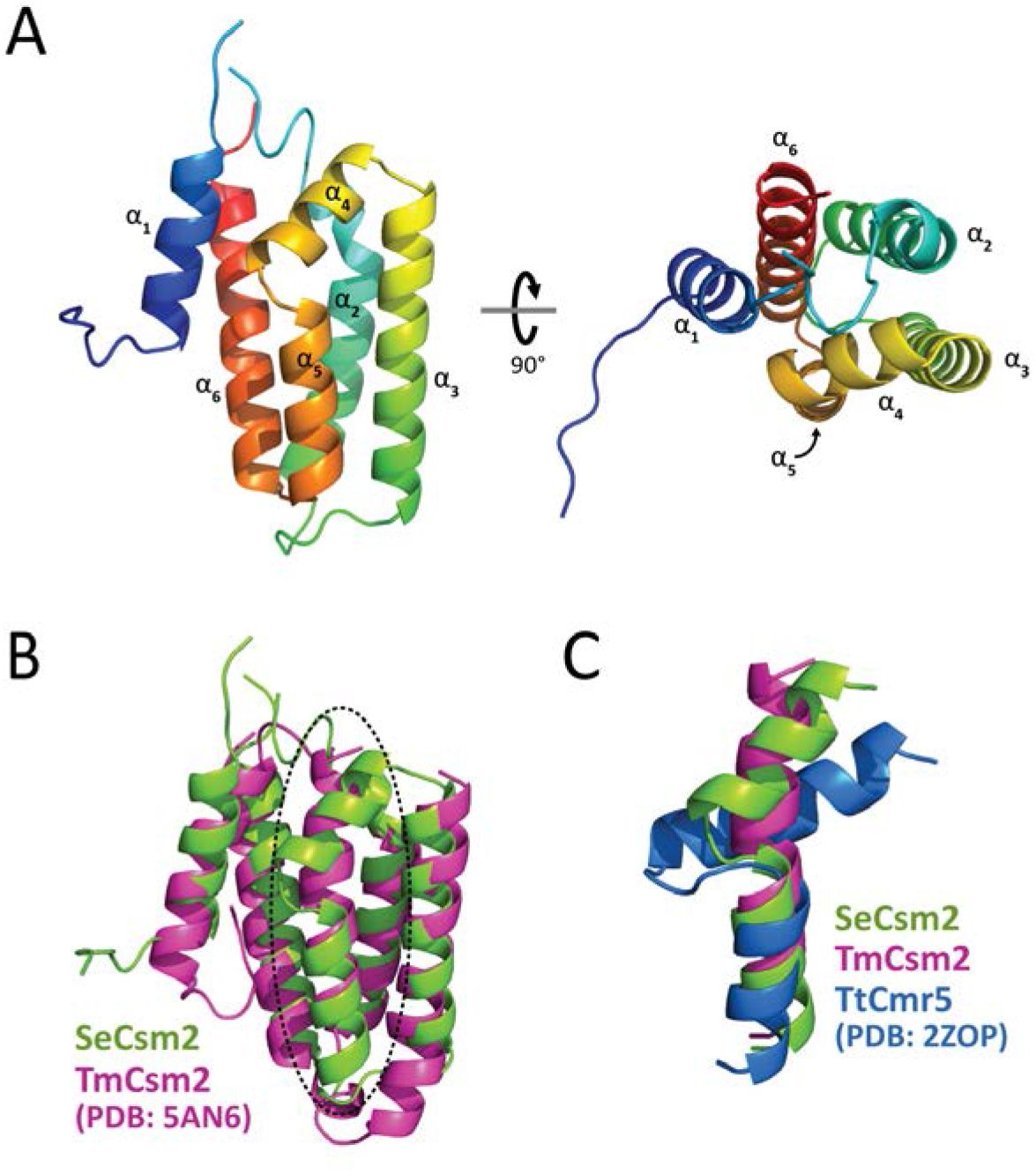
Molecular Architecture of *S. epidermidis* Csm2. (A) Two perpendicular views of the crystal structure of SeCsm2 show an exclusively α-helical fold consisting of 6 helices. Csm2 is colored in a rainbow from the N- (blue) to C- (red) termini. A portion of the N-terminal affinity tag was present in the density and is included in the structure. (B) Structural alignment of SeCsm2 (green) and the putative monomeric version of TmCsm2 (Gallo et al., 2016) (magenta). The overall architecture is similar with a root-means-squared-deviation (RMSD) between structures of 1.98 Å. One significant difference is the rearrangement of TmCsm2 α_3_ to form two helices (α_4_-α_5_) in SeCsm2 (dashed oval). (**C**) Structural alignment of SeCsm2 α_4_-α_5_ (green) with Type III-A TmCsm2 α_3_ (magenta) and Type III-B TtCmr5 α_4_-α_5_ (Sakamoto et al., 2010) (blue). The helix architecture of Type III-A and Type III-B Csm2 homologs reveals an intermediate organization in SeCsm2. (A)

SeCsm2 shows only two small positively charged electrostatic surfaces that could indicate a nucleic acid binding region (**Figure 2 - Figure supplement 1F**). In contrast, TtCmr5 exhibits a large positively charged electrostatic surface that has been implicated in target nucleic acid binding (Venclovas, 2016). When SeCsm2 and TtCmr5 are aligned, the positive patches are situated in different regions of the molecules, which may suggest alternate structural organizations of effector complex minor filaments. Additionally, a positively charged surface in the TmCsm2 dimer has been observed (Gallo et al., 2016), although it is not clear that the dimer represents a biologically relevant conformation. The structure of SeCsm2 or the packing in the crystal do not suggest a nucleic acid binding mode either, making the positively charged Loop 1 in SeCsm2 a likely candidate for a loop involved in nucleic acid binding. Models for TmCsm2 and TtCmr5 forming oligomers suitable for binding to RNA have been proposed (Venclovas, 2016) and a similar model could be built using SeCsm2.

### *S. epidermidis* Csm3 represents a minimized CRISPR backbone protein

Purification of nucleic acid-free *S. epidermidis* Csm3 (SeCsm3) was challenging due to its inherent ability to bind non-specifically to nucleic acids. Apparently pure SeCsm3 was contaminated with bound nucleic acids, which formed corkscrew-like aggregates of various lengths when visualized by negative stain electron microscopy (**Figure 3 - Figure supplement 1A**). Nucleic acid-free SeCsm3 was only produced after very slow loading onto a strong cation exchange resin and collecting the flow through of the column, which contained pure SeCsm3 (**Figure 3 - Figure supplements 1B-C**). Crystallization trials were performed on both nucleic acid-free and -bound SeCsm3, however only the nucleic acid-free protein produced high resolution diffracting crystals (**Figure 3 - Figure supplement 1D**). Although data were collected on both native and selenomethionine (SeMet) substituted crystals, only the SeMet crystals were used in the final structure determination due to the crystals diffracting to higher resolution. Phasing using SeMet was not possible due to the weak anomalous signal, later explained by the location of many of the SeMet in disordered regions of the structure (75% in molecule A and 50% in molecule B) (**Figure 3 - Figure supplement 1E**). No cysteines are present in SeCsm3, eliminating the use of selenocysteine as an alternative labeling strategy. Structure determination was ultimately accomplished by experimental SAD phasing from SeMet-SeCsm3 crystals derivatized with samarium (III) chloride. The unit cell contains two molecules in the asymmetric unit (**Figure 3 - Figure supplement 2A**). The final model of the SeCsm3 dimer (R_work_/R_free_ = 23.0%/26.7%) was obtained from data to 2.40 Å and consists of residues 1 to 21, 31 to 65, 74 to 124, and 137 to 214 (**Figure 3 - Figure supplement 1E** and **Supplementary Table 2**). The structure is identical to a recently published crystal structure of SeCsm3 (Y. Q. Zhao et al., 2018). Experiments combining size-exclusion chromatography with multi-angle light scattering analysis (SEC-MALS) show a molecular weight consistent with a monomer of SeCsm3 (data not shown), confirming that the dimer in the crystal is a result of crystal packing and does not reflect the oligomerization state in solution.

SeCsm3 consists of three distinct domains: 1) an RRM fold, 2) a β-hairpin, and 3) an isolated α-helix (**Figure 3A**). The RRM in SeCsm3 contains five anti-parallel β-strands and four α-helices (β_1_α_1_α_2_β_2_β_5_α_4_α_5_β_6_β_7_) with an insertion in the middle, between β_2_ and β_5_, to form the β-hairpin (β_4_ and β_5_) and the additional helix (α_3_) (**Figure 3B**). As expected for a molecule that binds nucleic acids, the electrostatic potential map shows a highly positively charged region spanning α_1_, α_5_, β_2_, β_3_, β_4_, and the loops surrounding them (**Figure 3 - Figure supplement 2B**). The crystallographic dimer shows the positive patches of each monomer facing one another, partially shielding them from the solvent (**Figure 3 - Figure supplement 2C**). The monomers of the crystallographic dimer are identical, with a relative-mean-standard-deviation (RMSD) of 0.15 Å. To ascertain the importance of conserved regions in the protein, the sequence of SeCsm3 was compared against 150 homologous sequences using the ConSurf server (Ashkenazy, Erez, Martz, Pupko, & Ben-Tal, 2010; Landau et al., 2005) and the conserved regions were analyzed. The results show an abundance of conserved residues in the positive patch of the protein and low conservation in other regions (**Figure 3 - Figure supplement 2D**), providing further evidence that the positively charged surface corresponds to a region of functional importance.

**Figure 3.**
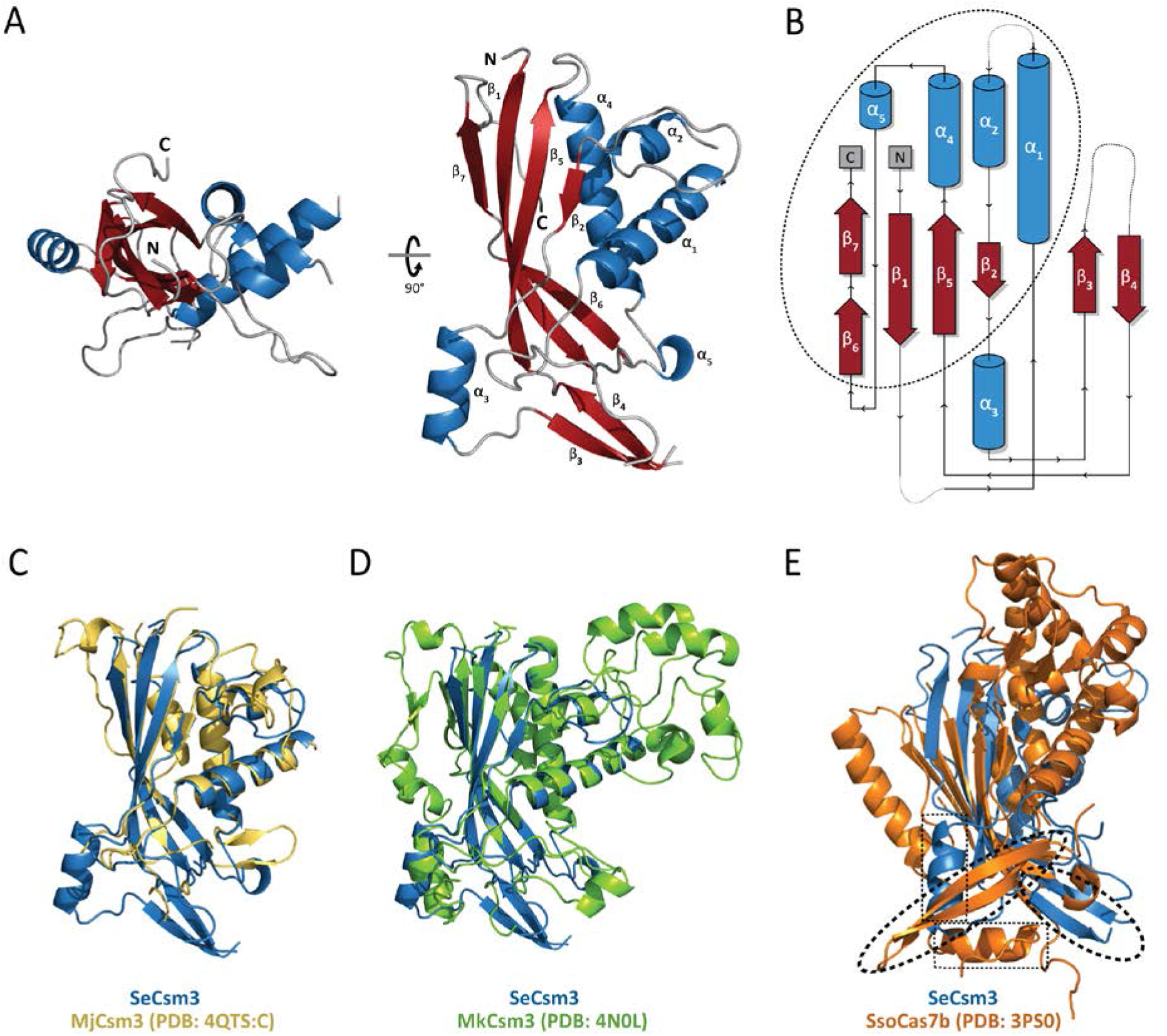
Molecular Architecture of *S. epidermidis* Csm3. (**A**) Two perpendicular views of the crystal structure of SeCsm3 show that it is composed of an RRM fold consisting of a five-stranded anti-parallel β-sheet and four helices, a β-hairpin, and an isolated α-helix. Sheets are colored in red and helices in blue. (**B**) Topology diagram of SeCsm3 shows that the N- and C-termini sequences form the RRM fold (dashed oval), with the other domains located at the center of the polypeptide. (**C-E**) Structural alignment of Cas7 family proteins. SeCsm3 is aligned with (**C**) MjCsm3 (Numata et al., 2015) (RMSD of 1.68 Å over 134 C_α_ atoms), (**D**) MkCsm3 (Hrle et al., 2014) (RMSD of 1.98 Å over 153 C_α_ atoms), and (**E**) SsoCas7b (Lintner et al., 2011) (RMSD of 2.99 Å over 122 C_α_ atoms). The β-hairpin (dashed ovals) and helix (dashed rectangle) in SsoCas7b are rotated with respect to the ones in SeCsm3. The alignments show the wide variation in the backbone subunit architecture of effector complexes in different types and subtypes.

SeCsm3 is known to cleave target RNA (Samai et al., 2015) and mutating D32 abrogates this activity without impairing crRNA processing or complex formation (Samai et al., 2015), strongly suggesting that D32 is located in the RNA cleavage active site. Mapping of D32 in the structure shows that it is found just after a disordered loop and in the vicinity of many other highly conserved residues that have been shown to be important for modulating crRNA length (H18, D100, E120, K122, E124, and R129) (Hatoum-Aslan et al., 2013; Makarova et al., 2011). All these residues reside in the positive patch and many face in the same direction. Clustered together around D32 are H18, E124, R127, R137, R141, and R187. Based on sequence comparisons, H18 has been suggested to be part of the active site, although mutational studies of this residue in *S. epidermidis* have shown no effect on effector complex formation, crRNA binding, or target cleavage (Hatoum-Aslan et al., 2013). Similar mutational studies of this conserved residue in other Type III Csm3 homologs have shown varied results, with no effect shown on CRISPR function in *Streptococcus thermophilus* Csm3 (Tamulaitis et al., 2014) and loss of target RNA binding in *Methanopyrus kandleri* Csm3 (Hrle et al., 2013). This suggests that this basic residue has lost its function in some Csm3 homologs, but kept its functionality in others. The presence of acidic amino acids D32 and E124 suggests that they could be involved in cation binding. Finally, the presence of positively charged residues, such as K122 and R129 is also consistent with a region involved in binding nucleic acids. Overall, the clustering of H18, D32, E124, R127, R137, R141 and R187, all highly conserved amino acids, strongly suggests that these amino acids are part of the active site involved in target RNA cleavage. Importantly, D100, E120, and K122, which have been shown to play a role in the regulation of crRNA length are further away and not in the vicinity of the putative active site.

Multiple effector backbone subunit structures have been determined for Type III-A (Hrle et al., 2013; Numata, Inanaga, Sato, & Osawa, 2015), Type III-B (Benda et al., 2014; Osawa, Inanaga, Sato, & Numata, 2015; Ramia, Spilman, et al., 2014; Zhu & Ye, 2015), Type I-D (Hrle et al., 2014), Type I-E (Jackson et al., 2014; Lintner et al., 2011; H. Zhao et al., 2014), and Type I-F (Pausch et al., 2017) CRISPR systems, and each has shown the canonical RRM core region decorated with multiple, varying domains that may or may not play a role in its function. Type III-A Csm3 structures have been solved from *Methanocaldococcus jannashii* (MjCsm3; PDB: 4QTS:C,D)(Numata et al., 2015) and *Methanopyrus kandleri* (MkCsm3; PDB: 4N0L)(Hrle et al., 2013). Whereas the structures of SeCsm3 and MjCsm3 share the same core architecture of the RRM fold (**Figure 3C**), sequence identity is low, at only 38% identity. Unfortunately, in MjCsm3, the region corresponding to the β-hairpin and isolated α-helix in SeCsm3 is disordered, preventing structural comparison of these regions (**Figure 3C**). The main structural difference between SeCsm3 and MjCsm3 occurs at the C-terminus, where an insertion from E213 to E224 in MjCsm3 accounts for roughly half of the residues that form the interface with MjCsm4 (Numata et al., 2015). When these residues were mutated to alanine, the interaction between the two subunits was lost. As SeCsm3 does not contain this region, it suggests that an alternate interaction surface must be present between SeCsm3 and SeCsm4. In contrast to MjCsm3, the structural differences between SeCsm3 and MkCsm3 are evident (**Figure 3D**). MkCsm3 contains a large N-terminal helical zinc-binding domain and a C-terminal helical domain that are completely absent from SeCsm3 (**Figure 3D**). When comparing the electrostatic potential maps of SeCsm3 and MkCsm3, there is a dramatic shift in the orientation of the putative RNA binding surface. SeCsm3 has the binding surface at the center of the molecule, whereas the MkCsm3 binding surface is on a more proximal region, classified as the lid domain (Hrle et al., 2013). SeCsm3 also has a more occluded, channel-like binding region (**Figure 3 - Figure supplement 2B**) compared to the flattened binding region on MkCsm3 (Hrle et al., 2013). The variation of these binding region characteristics may change how successive Csm3 subunits interact to form the backbone of the effector complex and suggest that the structure of the effector complex may show differences even amongst members of the same subtype.

Comparing the SeCsm3 structure to effector complex backbone subunits from other types shows a more diverged RRM architecture as well as variation in domain organization. When SeCsm3 is aligned to *Sulfolobus solfataricus* Cas7b (SsoCas7b; PDB: 3PS0)(Lintner et al., 2011), the equivalent Type I-E backbone subunit, the most prominent feature is the reorganization of the β-hairpin (**Figure 3E**). The β-hairpin in SsoCas7b is flipped more than 90° with respect to the equivalent β-hairpin in SeCsm3, causing the isolated α-helix to be oriented almost perpendicular to the equivalent α-helix in SeCsm3 (**Figure 3E**). In addition, SsoCas7b α_2_ (corresponding to α_1_ in SeCsm3) is positioned such that the putative binding region of SeCsm3 would be occluded. These rearrangements relative to SeCsm3 create a more open binding surface, with the majority of the positively charged patch located around the β-hairpin (Lintner et al., 2011). In SsoCas7b, the additional helical domain proximal to the RRM fold as well as the C-terminal helical domain coincide surprisingly well to the respective C- and N-terminal helical domains found in MkCsm3. It is presently unknown how these rearrangements in SsoCas7b change the effector complex backbone quaternary architecture relative to the structure of the SeCsm3 major filament.

### *S. epidermidis* Cryo-EM Structure Elucidates the Quaternary Architecture of the Major Filament Subcomplex

In order to study the overall architecture of the Cas10-Csm effector complex, the complex was purified directly from *S. epidermidis* RP62a cells using a Cas10 subunit with a His tag attached. Purification of the *S. epidermidis* effector complex yielded a homogeneous sample where all five subunits were present and of the expected molecular weight (**Figure 4A**). The purified effector complex was characterized both by negative stain and cryo-EM. To prevent aggregation of the complex, which caused particles to overlap in the EM grid, the salt concentration was increased from 200 mM to 500 mM NaCl (**Figure 4B**). The high salt species still showed the presence of five subunits and isolation of the crRNA from the purified complex by phenol-chloroform extraction yielded two predominant species corresponding to pre-crRNA and mature crRNA, consistent with what was shown in previous studies (**Figure 4C**) (Hatoum-Aslan et al., 2013). Negative stain EM analysis was performed to generate an initial model for cryo-EM studies (data not shown), which showed a complex of the expected shape and size. Initial analysis of the cryo-EM data showed detailed 2D class averages, with most corresponding to views where the effector complex lies with the long dimension parallel to the ice layer (**Figure 4D**). The data showed the presence of only one particle species. A 3D density map was calculated from 107,620 particles using the negative stain model as a starting point. After post-processing steps were performed, including per-particle CTF estimation and particle polishing, the final volume had an overall resolution of 5.2 Å, as estimated by the gold standard FSC criterion (**Figure 5 - Figure supplement 1A**). Local resolution estimates (Kucukelbir, Sigworth, & Tagare, 2014) agreed with this resolution assignment (**Figure 5 - Figure supplement 1B**). The structure shows a slightly helical stem with an extended domain at the base of the molecule (**Figure 5A**). The helical stem was clearly of higher resolution than the extended domain, whose resolution was estimated at around 7-9 Å (**Figure 5 - Figure supplement 1B**).

**Figure 4.**
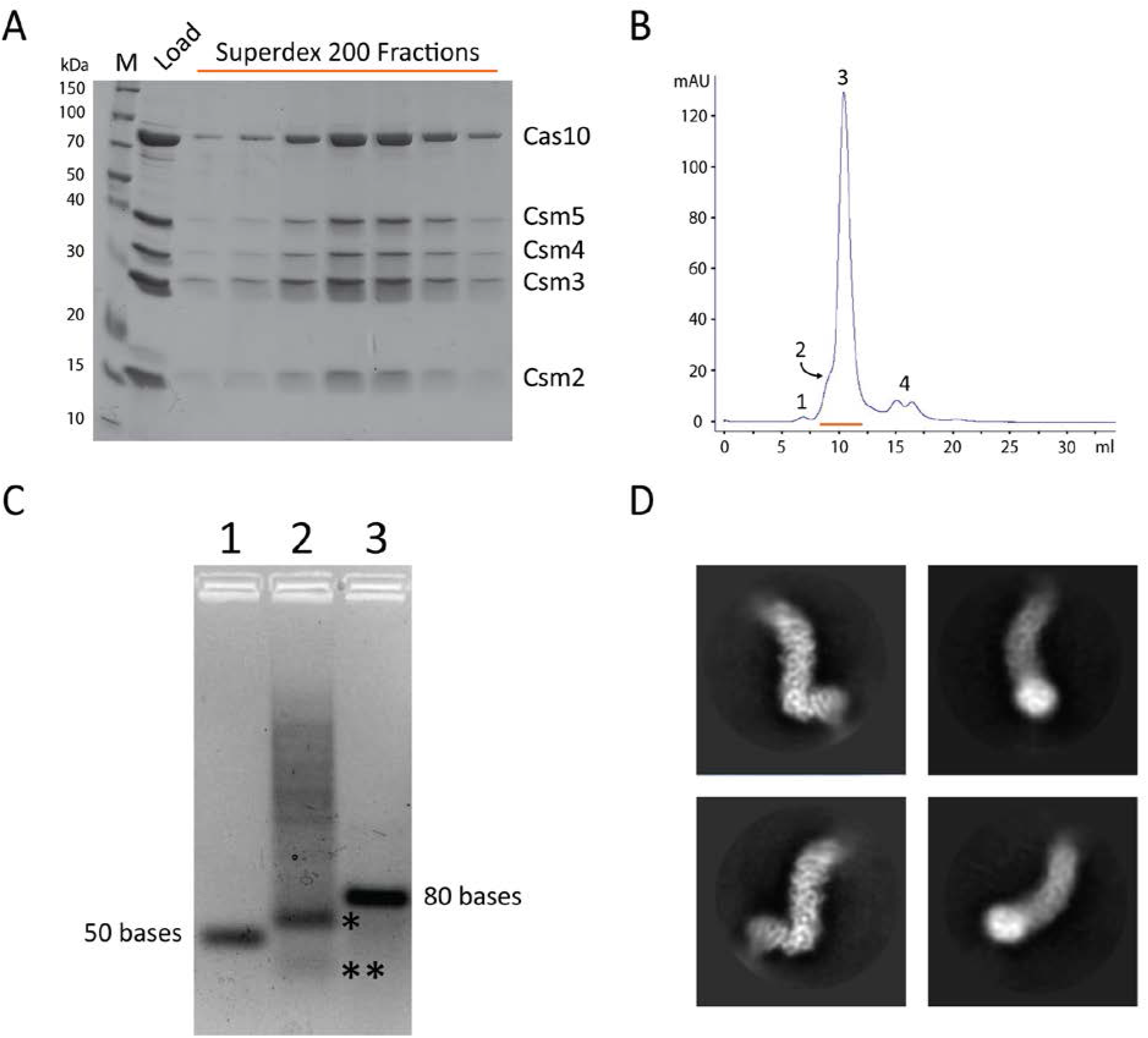
Purification of *S. epidermidis* CRISPR Effector Complex. (**A**) Coomassie blue stained 12% SDS-PAGE of the purified SeCas10-Csm effector complex. The gel shows fractions collected around the peak eluted from the column. All five proteins expected in the complex are present in the peak. The identity of the proteins is shown on the right. M, molecular weight markers. (**B**) Superdex 200 trace of the SeCas10-Csm complex purification. Indicated peaks correspond to the column void volume (1), SeCas10-Csm aggregate peak (2), SeCas10-Csm non-aggregate peak (3), and contaminants (4). The red line marks the elution fractions loaded into the gel shown in **A**. (**C**) crRNA isolated by phenol-chloroform extraction from the purified complex (lane 2) is present as both the 71nt pre-crRNA (*) and the mature crRNA (**) of undetermined length. Lane 1 is a 50 base DNA oligonucleotide marker and lane 3 is an 80 base DNA oligonucleotide marker. (**D**) Cryo-EM 2D class averages of the effector complex show well-defined features within the core of the molecule, but heterogeneity at the extremes.

**Figure 5.**
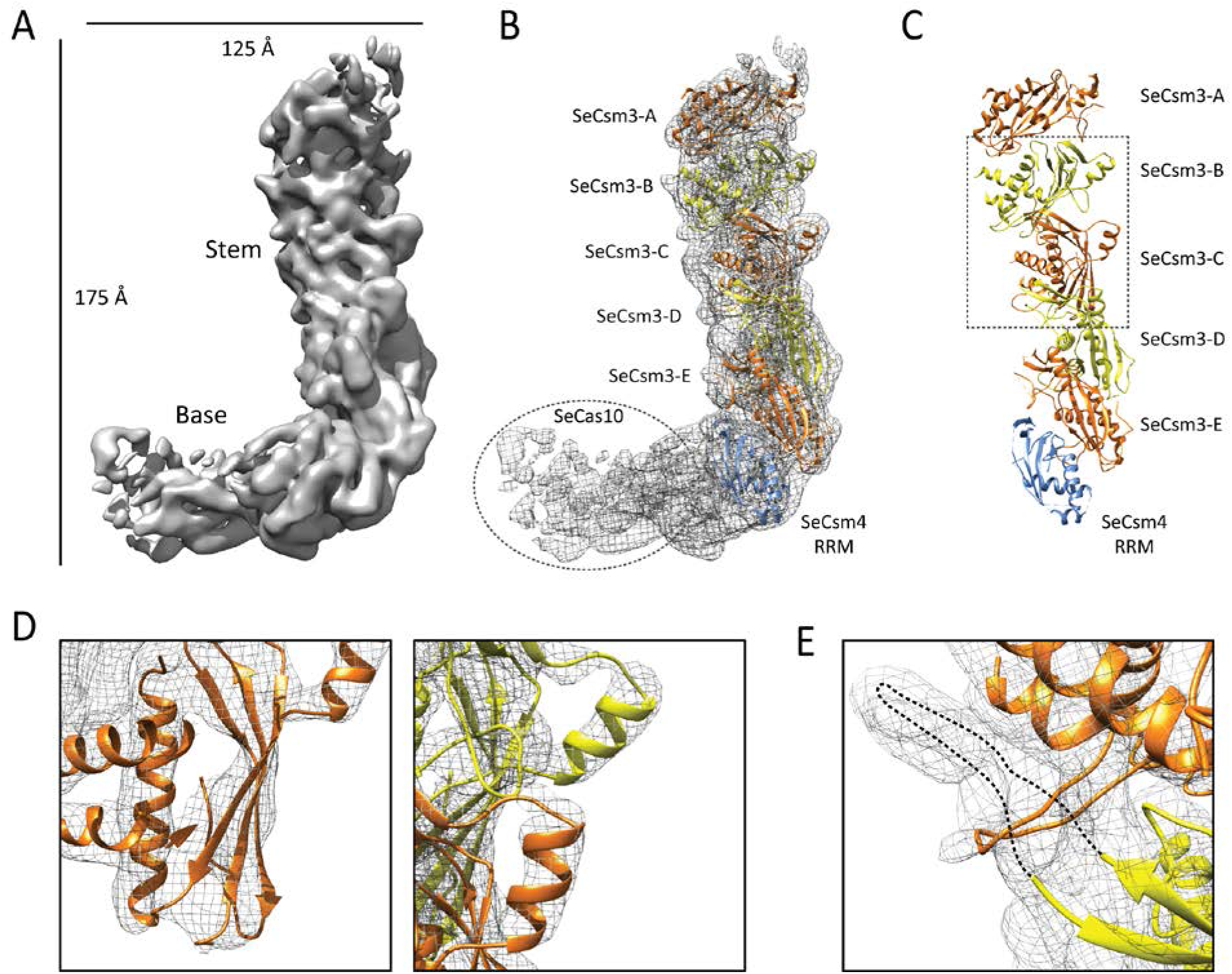
Overview of the Cryo-EM Structure of the Major Filament Subcomplex. (**A**) A 5.2 Å cryo-EM reconstruction of the SeCas10-Csm subcomplex shows a helical stem region adjoining the non-helical base region. The stem is formed by five SeCsm3 subunits while the base includes SeCsm4 and SeCas10. (**B**) SeCsm3 and a putative SeCsm4 RRM docked into the cryo-EM density map. Five SeCsm3 (alternately colored orange and yellow) and one SeCsm4 RRM (blue) were confidently placed in the density. Crystal structures of homologous structures or homology models for SeCas10 and SeCsm4 did not fit well into the density at the base. Additional density at the base was putatively assigned as SeCas10 (dashed oval). (**C**) The SeCsm3 crystal structures and the SeCsm4 RRM form a helical arrangement, with the RRM β-strands facing the outside of the helix and the β-hairpin on the inside of the helix. Regions in the dashed box are shown in **D** and **E**. (**D**) Close-up views of the SeCsm3 secondary structure elements show good agreement with the cryo-EM density. (**E**) Disordered regions from the SeCsm3 crystal structure show density in the cryo-EM map. These regions of density were putatively assigned as an extension of each SeCsm3 β-hairpin (dashed lines).

In order to locate the positions of the SeCsm3 crystal structure in the cryo-EM map, a 6 dimensional search using Essens (Kleywegt & Jones, 1997) was done. The search identified five separate SeCsm3 locations, in addition to an RRM fold associated at the base of the backbone stem, which was assigned as SeCsm4 (**Figures 5B** and **5C**). A similar search with the crystal structure of SeCsm2 did not yield any solutions, consistent with visual inspection of the volume, and which suggests that SeCsm2 is not found in this complex. The SeCsm3 subunits accounted for most of the density in the stem, whereas the RRM fold assigned to SeCsm4 filled a small region at the base of the stem. No density corresponding to SeCsm5, which is expected to be located at one end of the stem (**Figure 1B**) was identified in the structure. The remaining density at the base of the stem was generally assigned to SeCsm4 and SeCas10, which have been shown in other structures to interact directly (Y. Shao et al., 2013; R. H. Staals et al., 2014; Taylor et al., 2015; Wiedenheft et al., 2011) and which can be purified recombinantly as a binary complex (data not shown). The density at either end of the complex is of lower resolution and secondary structure elements are harder to discern. This could potentially be due to slightly different relative orientations between the base and stem. To address this, multi-body refinement of the base and stem as two separate bodies was performed (**Figure 5 - Figure supplement 1C**). However, this did not improve the quality of the map (**Figure 5 - Figure supplement 1B**).

The SeCsm3 crystal structure fits well at five locations (labeled as SeCsm3-A to E) (**Figure 5B**). Although the model accounts for most of the density around these subunits, the map shows additional density for regions that are disordered in the crystal structure, such as an extension of the incomplete β-hairpin (**Figure 5E**). Flexible fitting of the structure using MDFF (Trabuco, Villa, Mitra, Frank, & Schulten, 2008) into the cryo-EM map showed that the crystal structure fits without major changes, aside from slight re-orientations of some of the surface loops (**Figure 5 - Figure supplement 1D**). SeCsm3-A has the worst density, but overall it fits well. Nevertheless, we cannot completely rule out that the density assigned to SeCsm3-A corresponds to SeCsm5, especially if the two subunits are structurally very similar. Homology models for SeCsm4 and SeCas10 were generated, but did not fit the density adequately and were therefore not included in the final structural model. Nevertheless, the homology models served to show that the density at the base of the stem was of the shape and size to accommodate both SeCas10 and SeCsm4. A partial RRM model for SeCsm4 was built using SeCsm3 as a template. The SeCsm4 RRM lies next to SeCsm3-E in the stem and makes contacts with it (**Figure 5C**). Superposition of the MjCsm3 subunit in the MjCsm3-MjCsm4 complex (Numata et al., 2015) on SeCsm3-E showed that SeCsm4 is positioned similarly to MjCsm4, confirming the assignment of the density to the SeCsm4 subunit. Overall, it was possible to assign density for five SeCsm3 subunits, one SeCsm4 subunit and one SeCas10 subunit (**Figure 6A**). The SeCsm2 and SeCsm5 subunits are not present in the cryo-EM structure.

**Figure 6.**
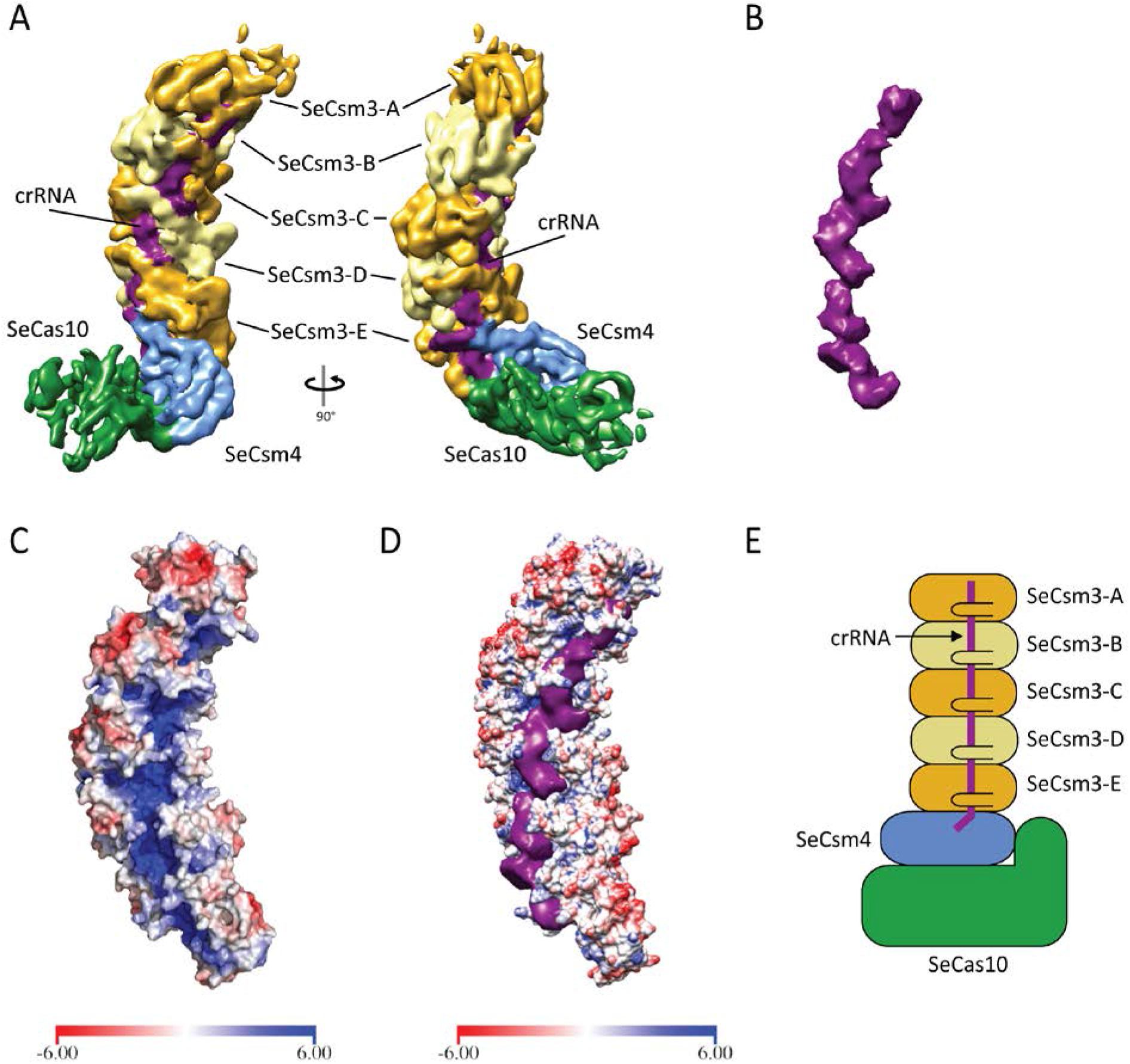
The Effector Subcomplex Shows the crRNA in a Conserved Positively Charged Channel. (**A**) Two orthogonal views of the cryo-EM map colored by subunit identity. Cryo-EM density was assigned to each subunit in the complex after crystal structure docking. Density is assigned for five SeCsm3 (gold and yellow), one SeCsm4 (blue), and one SeCas10 (green), with additional density within each SeCsm3 putatively assigned to crRNA (purple). (**B**) Putative crRNA density isolated from the SeCas10-Csm subcomplex shown in **A**. The density follows a continuous path along the length of the stem. (**C**) The SeCsm3 stem is shown in accessible surface representation colored by electrostatic potential calculated using APBS (Baker et al., 2001). A highly positively charged channel (blue) is seen traversing the length of the stem. (**D**) Putative crRNA density (purple) overlaid on the electrostatic potential map of SeCsm3 stem. The positively charged channel surrounds the negatively charged crRNA, providing further evidence that the density corresponds to the crRNA. The bars at the bottom of **C** and **D** map the electrostatic potential values from -6 k_B_T/e_c_ [red] to 6 k_B_T/e_c_ [blue]. (**E**) Schematic diagram showing the composition of the SeCas10-Csm subcomplex. Csm2 and Csm5 are not present in the subcomplex.

After assigning density to each of the three subunits, a continuous region of unaccounted for density was located within each SeCsm3 subunit (**Figure 6B**). This extra density was putatively assigned as the crRNA due to its shape and location (**Figure 6A**). To support the assignment, an electrostatic potential map was calculated for the docked SeCsm3 crystal structures forming the stem. A highly positively charged channel was observed traversing the length of the stem (**Figure 6C**). When the putative crRNA density was aligned with the electrostatic potential map, the density fit remarkably well into the positively charged channel (**Figure 6D**), allowing us to confidently assign this density to the crRNA. Taken together, the identification suggests that the structure corresponds to an effector subcomplex with a stoichiometry of SeCsm3_5_SeCsm4_1_SeCas10_1_crRNA_1_ (**Figures 6A** and **6E**). The crRNA is of undetermined length, but given the continuous density and the fact that each SeCsm3 binds 6 nucleotides (Hatoum-Aslan et al., 2013) its length is at least 30 nucleotides long, but probably slightly longer.

The overall architecture of the subcomplex (**Figures 6A** and **6E**) is similar to that of other CRISPR-Cas complexes and in agreement with the predicted overall arrangement (**Figure 1B**). The stem is formed by five SeCsm3 subunits with the SeCas10/SeCsm4 subunits at the base of the complex and almost perpendicular to the stem (**Figure 6A**). SeCsm4 makes clear contacts with SeCsm3-E and the crRNA seems to extend into SeCsm4. SeCas10 also appears to contact SeCsm3-E, but this is not as clear as it is not possible to assign unambiguously density to SeCsm4 and SeCas10. The crRNA runs along the positively charged groove formed by SeCsm3-A to E (**Figure 6D**) and appears to extend into the RRM domain of SeCsm4 (**Figure 6A**). It is not clear whether it continues into SeCas10. SeCsm5 is expected to reside on the other end of the stem next to SeCsm3-A (**Figure 1B**), but it is not possible to assign density for it.

The availability of an atomic model for the SeCsm3 stem helps locate many of the residues that have been implicated in different biochemical activities, such as forming the SeCsm3-SeCsm3 interface (Walker, Chou-Zheng, Dunkle, & Hatoum-Aslan, 2017), crRNA binding and maturation (Hatoum-Aslan, Maniv, Samai, & Marraffini, 2014; Hatoum-Aslan et al., 2013; Walker et al., 2017), target RNA binding and cleavage (Hatoum-Aslan et al., 2013; Samai et al., 2015; Walker et al., 2017), and protein stability (Hatoum-Aslan et al., 2014). Biochemical and functional studies suggest that residues H18 and D32 are involved in RNA target cleavage, residues K52, K54, R56, D100, E120, E124 and E140 affect crRNA maturation, amino acids G182, G183, and G185 have an impact on protein stability, while residues K4 and D179 form part of the SeCsm3-SeCsm3 interface (**Supplementary Table 3** and **Figure 6 - Figure supplements 1A-B**). Many of these residues are amongst the most highly conserved ones in the SeCsm3 backbone (**Figure 6 - Figure supplement 1C**) and in general can be found in or near the highly positively charged channel (**Figure 6C**), suggesting that their role is not confined to SeCsm3, but extends to all members of the Type III-A and possibly Type III-B subtypes. Analysis of both free SeCsm3 and SeCsm3 in the effector complex show that the four sets of residues are spatially separated from one another (**Figure 6 - Figure supplements 1A-B**). As mentioned above, many highly conserved residues involved in target cleavage (Hatoum-Aslan et al., 2013) cluster around H18 and D32 and form the putative active site in SeCsm3. The SeCsm3-SeCsm3 interface is extensive and the two residues biochemically identified as part of the interface (Walker et al., 2017) are facing each other in adjacent subunits, suggesting that K4 and D179 form a salt bridge that helps stabilize the stem in the effector complex. Residues involved in crRNA binding and maturation all cluster in the same region (**Figure 6 - Figure supplement 1B**). Three basic amino acids, K52, K54 and R56 face the region where the crRNA is located in the effector complex (**Figure 6 - Figure supplement 2A**). D100 is near a highly conserved lysine, K122, in the adjacent subunit and it is possible that they form an inter-subunit ion pair. E140 is further away and makes an intra-subunit salt bridge with R37 (**Figure 6 - Figure supplement 1B**). Although R37 is not strictly conserved, it is either an arginine or a lysine, suggesting that the salt bridge is conserved and may play a role in stabilizing the tertiary structure of SeCsm3. E124 is also found near the interface between two subunits, but not the same subunits as for E140 and D100. Finally, the three glycines are part of a loop formed by four consecutive glycines and facing the crRNA. Overall, it is possible to classify the mutated amino acids into three large groups, those involved in making contacts at the interface between adjacent subunits, those forming part of the putative active site, and those involved in structural stability (**Figure 6 - Figure supplements 1A-B**).

The cRNA runs along the length of the SeCsm3 stem and travels near many conserved amino acids. Not surprisingly, the active site residues are not found abutting the crRNA density (**Figure 6 - Figure supplement 2A**). Instead, the putative active site is found adjacent to the crRNA and facing a cavity that could easily accommodate the target RNA strand (**Figure 6 - Figure supplement 2B**).

## Discussion

The structural studies presented here describe high-resolution crystal structures of SeCsm2 and SeCsm3 and a medium resolution cryo-EM structure of a Cas10-Csm subcomplex that shows the quaternary architecture of the major filament of the complex. The structure of SeCsm2 shows a small five-helix bundle protein similar to the orthologous Cmr5 found in Type III-B CRISPR systems (Sakamoto et al., 2010), as well as to other Type III-A Csm2 proteins (Gallo et al., 2016; Venclovas, 2016). However, the SeCsm2 structure shows important differences between both Type III-A and III-B Csm2 orthologs, such as the presence of a small helix that breaks a longer helix. This architecture shows SeCsm2 has some of the characteristics of other Type III-A Csm2 proteins, but also contains elements found in the Type III-B Cmr5 proteins. There are currently no known functions ascribed specifically to SeCsm2. However, other complexes have described orthologs acting to stabilize the binding of target DNA (Hayes et al., 2016; H. Zhao et al., 2014) by forming a second stem parallel to the SeCsm3 stem (Kazlauskiene, 2016) (**Figure 1B**). Knockout studies have shown that SeCsm2 is one of the components required for crRNA maturation, but mutational studies do not implicate any of the conserved residues in this maturation role (Walker et al., 2017). Although the crystal structure helps understand the organization of SeCsm2, its absence in the subcomplex described here precludes any further identification of its role in the effector complex. In addition, it is not clear whether the differences observed between different Type III-A Csm2 proteins and their Type III-B orthologs may cause differences in the overall conformation of the effector complex.

In contrast to SeCsm2, the crystal structure for SeCsm3 provides important information on the structure and function of the subunit and how it relates to its structural and functional roles within the effector complex. SeCsm3 represents a minimized version of proteins that form the backbone in CRISPR-Cas effector complexes. Unlike other Csm3 proteins and Csm3 orthologs, SeCsm3 is formed primarily by the core RRM fold and a β-hairpin. The SeCms3 crystal structure shows the position of many important and highly conserved amino acids as well as the position of the putative active site responsible for RNA target cleavage suggested by mutagenesis studies to be located in the vicinity of D32 (Samai et al., 2015). Aside from H18 and D32, the putative active site contains many residues normally associated with active sites of proteins involved in nucleic acid cleavage, such as arginines, lysines, and additional acidic residues. However, unlike other ribonucleases, such as ribonucleases H1 and H2 (Tadokoro & Kanaya, 2009), there is no clustering of acidic residues potentially involved in cation binding. Other endoribonucleases in CRISPR systems, such as *T. thermophilus* Cse3 (PDB: 2Y8W) (Sashital, Jinek, & Doudna, 2011) and *Methanococcus maripaludis* Cas6b (PDB: 4Z7K) (Y. M. Shao et al., 2016), show different active site locations when aligned to SeCsm3, suggesting a different mechanism of cleavage for events associated with crRNA maturation and target RNA degradation. Determining whether SeCsm3 uses a two metal cleavage mechanism or other common ribonuclease catalytic strategy remains to be elucidated and may require structures of complexes with target RNA.

The cryo-EM structure shows a subcomplex formed by three of the subunits, SeCsm3, SeCsm4, and SeCas10 (**Figure 6E**). Neither SeCsm2 nor SeCsm5 could be located in the subcomplex. The absence of these subunits is intriguing as the purified complex showed a single migrating species in size-exclusion chromatography (**Figure 4B**) with all five protein components present (**Figure 4A**) as well as the crRNA (**Figure 4C**). One possible explanation for the loss of the subunits is the high salt concentration used to mitigate complex aggregation. SeCsm2 and SeCsm5 may also associate weakly in the complex without the presence of the target nucleic acid, enabling the vitrification process utilized in cryo-EM sample preparation to pull apart these components from the complex. Previous studies have shown that complex formation requires SeCsm3, SeCsm4, and SeCas10, while a subcomplex of these components forms when SeCsm2 or SeCsm5 are knocked out (Hatoum-Aslan et al., 2014). These subcomplexes were shown to include only the intermediate 71 nucleotide pre-crRNA (Hatoum-Aslan et al., 2014), while crRNA isolated from the cryo-EM complex shows both intermediate and mature crRNA (**Figure 4C**). This further supports the notion that the purified complex was intact, active, and containing all subunits, as it was capable of processing pre-crRNA to the correct mature length, but lost some subunits during the vitrification process. The subcomplex in the cryo-EM structure thus represents a partial complex formed by loss of some of the subunits after formation, and not a partially assembled complex.

The assignment of different subunits to the density in the cryo-EM map was aided by the crystal structure of SeCsm3 as well as the presence of an RRM fold in SeCsm4. Prior studies had determined that SeCsm3 is required for proper effector complex assembly (Hatoum-Aslan et al., 2013), with multiple copies of Csm3 forming the backbone of Type III-A complexes (R. H. Staals et al., 2014; Tamulaitis et al., 2017). The structure confirms this and shows that SeCsm3 does form the backbone of the effector complex and that there are five SeCsm3 copies present (**Figures 5B** and **5C**). Although SeCsm5 is expected to cap one of the ends of the effector complex, the assignment of the density at the end of the stem as SeCsm3-A, rather than SeCsm5, was based on the presence of an RRM fold that fits well into the map, density that accounts for the entirety of SeCsm3, the existence of crRNA density within this subunit, and comparison with the cryo-EM structure of the only other Type III-A effector complex structure known, that of the *T. thermophilus* CRISPR Cas10-Csm complex (R. H. Staals et al., 2014), which shows six subunits in the backbone stem (**Figure 6 - Figure supplement 3A**). At the other end of the stem, the location of SeCsm4 was aided by the presence of an RRM fold in the density. Although SeCsm3 has a similar RRM fold, the density did not fit the SeCsm3 subunit well as there were clear differences in some regions described by either additional or missing density. In support of this assignment, the structure of a Csm3-Csm4 complex from *M. jannaschii* showed a similar interaction between the two proteins (Numata et al., 2015). Superposition of MjCsm3 on SeCsm3 in the complex positioned MjCsm4 in the same position as SeCsm4. Finally, the position of SeCas10 was deduced from the size and shape of the unaccounted for density, as well as comparisons of other orthologous Type III-B structures published describing the association of Cas10 (Cmr2) and Csm4 (Cmr3) (Jung et al., 2015; Y. Shao et al., 2013). Given the resolution of the cryo-EM map, the boundaries between the SeCsm3, SeCsm4 and SeCas10 are not well defined, but the general organization is.

The resolution of the cryo-EM map is uneven; the SeCsm3 stem is the most ordered region, while the two ends are less ordered, particularly the SeCas10 region (**Figure 5 - Figure supplement 1B**). This observation suggests that SeCas10 is either moving or does not have a tightly held position with respect to the SeCsm3 stem. It is not clear if this difference in relative orientation is due to the absence of SeCsm2, which could play a role in stabilizing the interaction, or simply reflects the flexibility of the complex. It is possible that the conformation of SeCas10 is not rigid and thus is part of its function. During target DNA binding, SeCas10 may undergo a conformational change to help capture and orient target DNA properly in the active site.

The position of the crRNA in the complex follows the expected path, winding its way along the positively charged and highly conserved groove created by the SeCsm3 subunits. Previously it was found that each SeCsm3 subunit binds six nucleotides (Hatoum-Aslan et al., 2013). The structure shows that the distance between identical residues in adjacent subunits is around 25 Å (**Figure 6 - Figure supplement 2B**), which is consistent with the distance spanned by six nucleotides in an extended conformation and giving a length of over 30 nucleotides for the crRNA in the complex structure. It appears that the crRNA phosphate backbone is protected by the protein, where the β-hairpin loop perpendicularly transverses the crRNA while the bases of the crRNA are exposed to the solvent, where it could easily interact with complementary target DNA or RNA. Residues that have been implicated in crRNA binding (Hatoum-Aslan et al., 2014; Hatoum-Aslan et al., 2013; Walker et al., 2017) are located in the vicinity (**Figure 6 - Figure supplement 2A**). The active site location is adjacent to the crRNA and exposed and, as expected, the crRNA does not come close to the putative SeCsm3 active site (**Figure 6 - Figure supplement 2A**). It is possible that once the crRNA recognizes and binds its target, the target strand enters the SeCsm3 active site where it can be cleaved if it is RNA. In the structure, there appears to be space to accommodate the target strand (**Figure 6 - Figure supplement 2B**), although it is possible that the stem has to move in a concerted fashion, as has been observed for other CRISPR-Cas effector complexes (Taylor et al., 2015; H. Zhao et al., 2014).

The only other Type III-A effector complex structure known is a low resolution structure from *T. thermophilus* (R. H. Staals et al., 2014), an organism which harbors various types and subtypes of CRISPR systems, including both Type III-A and III-B CRISPR subtypes. The Type III-A *T. thermophilus* complex comprises a full complex with a Cas101Csm23Csm36Csm42Csm51crRNA1 stoichiometry. Aligning the two structures shows similar major filament architectures (**Figure 6 - Figure supplement 3A**) even though the SeCas10-Csm structure represents a sub-complex missing SeCsm2 and SeCsm5 (**Figure 6E**). Superposition of the density maps from both structures show that the *S. epidermidis* subcomplex has a very similar overall structure to the Type III-A *T. thermophilus* complex (**Figure 6 - Figure supplement 3A**), with one major difference being the number of Csm3 and Csm4 subunits. The higher resolution cryo-EM structure of the SeCas10-Csm complex allows unambiguous assignment of the crystal structure of SeCsm3 to the cryo-EM map, where it is clear that there are only five SeCsm3 subunits. In addition, the volume also suggests the presence of only one SeCsm4 subunit. The superposition of the maps shows a similar size volume for the region assigned to one SeCsm4 and one SeCas10 as well as additional density to place one SeCsm5 at the other end of the stem. The *T. thermophilus* structure also suggests a path for the SeCsm2 subunits, forming a helical stem running parallel to the SeCsm3 stem. Overall, the comparison shows that the two Type III-A complexes are remarkably similar, even though the subunit stoichiometry may be different.

The structure of the *T. thermophilus* Type III-B complex is also known (Taylor et al., 2015). The Type III-B effector complex has a Cmr11Cmr21Cmr31Cmr44Cmr53Cmr61crRNA1 stoichiometry (the corresponding orthologs are Cmr1:Csm3, Cmr2:Cas10, Cmr3:Csm4, Cmr4:Csm3, Cmr5:Csm2, Cmr6:Csm5). Type III-B systems are an evolutionary offshoot from the Type III-A systems, where a Cas7 family gene duplication occurred. This led to the incorporation of three Cas7 family proteins (Cmr1, Cmr4, and Cmr6) into Type III-B effector complexes, whereas only two Cas7 family proteins (Csm3 and Csm5) are found in Type III-A effector complexes (Makarova et al., 2011). The two subtypes have the same effector complex stoichiometry when compared by gene family, Cas101Cas51Cas76Cas113crRNA1, where the Cas5 family includes Csm4/Cmr3 and the Cas11 family includes Csm2/Cmr5. However, when the structures are compared, it is clear that they are very different in overall architecture (**Figure 6 - Figure supplement 3B**). While the *S. epidermidis* effector complex has a boot-shaped overall architecture, the *T. thermophilus* Type III-B complex is much more rod-shaped and without a marked protrusion. Both complexes have in common the presence of a helical stem and a similar subunit arrangement, but overall the two complexes are quite different. The same is true when the *S. epidermidis* complex is compared to other types, such as the Type I-E represented by the *Escherichia coli* CASCADE complex (Mulepati, Heroux, & Bailey, 2014). The *E. coli* complex is also elongated and with a ‘seahorse’ architecture (Mulepati et al., 2014), but without any similarity in overall architecture or subunit stoichiometry. The main commonality amongst all these complexes is the presence of a helical arrangement of subunits to bind and present the crRNA. Overall, the diversity in subunit composition of the different types and subtypes is reflected in a wide array of effector complex structures, with the major commonality occurring in the arrangement of the subunits that bind the crRNA.

The structures presented help to understand better the overall architecture of the Type III-A Cas10-Csm effector complex. The structures help to identify the location and organization of functionally relevant residues identified by biochemical experiments, to understand the way that the Cas10-Csm complex binds the crRNA, and to detail the organization of the RNA cleavage site. This new structural information provides a structural foundation to determine the molecular mechanisms responsible for the function of the Type III-A CRISPR system.

## Materials and Methods

### SeCsm2 and SeCsm3 Recombinant Plasmid Construction and Expression

The coding region for the *csm2* and *csm3* genes were PCR-amplified from extracted *S. epidermidis* RP62a genomic DNA using the PCR primers for *csm2* (Csm2-F & Csm2-R) and *csm3* (Csm3-F & Csm3-R) (**Supplementary Table 4**). PCR-amplified genes were inserted into the pET21-derived expression plasmid pMCSG7 (Stols et al., 2007) through the ligation-independent cloning (LIC) region. Utilizing the LIC region introduces an in-frame 6X Histidine tag and a tobacco etch virus (TEV) cleavage site at the N-terminus of the protein. The constructs were confirmed by DNA sequencing (ACGT Inc, Germantown, MD) and subsequently transformed into Rosetta competent cells (EMD Millipore) for protein expression and purification. Both constructs were grown in Terrific Broth (TB) media supplemented with 100 μg/ml ampicillin and 34 μg/ml chloramphenicol at 37°C to an OD600 of 0.8. Cells were cooled on ice for 1 hour to reduce leaky gene expression before induction with 0.5 mM isopropyl β-D-1-thiogalactopyranoside (IPTG) for 18 hours at 16°C. Cells were harvested by centrifugation at 5,500 X *g* for 15 minutes at 4°C. Cell paste was then flash frozen in liquid nitrogen and stored at -80°C until purification.

Expression of selenomethionine substituted SeCsm2 and SeCsm3 was performed by growth of a 5 ml pre-culture in Luria Broth (LB) rich media grown for 8 hours at 37°C. A 10 ml M9 minimal media starter culture was inoculated with 1:1000 pre-culture and grown overnight at 37°C. One liter of M9 minimal media cultures were inoculated with 1:100 M9 starter and grown to an OD600 of 0.8. Cells were cooled on ice for 45 minutes before adding 100 mg lysine, 100 mg threonine, 100 mg phenylalanine, 50 mg leucine, 50 mg isoleucine, 50 mg valine, and 50 mg L (+) selenomethionine (ACROS Organics). After 15 minutes, cells were induced with 0.5 mM IPTG for 18 hours at 16°C. Cells were harvested and frozen following the same protocol as for the native proteins.

### Recombinant SeCsm2 Protein Purification

Pellets were thawed on ice for 30 minutes before adding 1:1 volume of resuspension buffer (100 mM Tris pH 8, 1M NaCl, 20 mM imidazole). Resuspended cells were incubated with 50 μM lysozyme, 0.12% Brij 58, 1 mM phenylmethylsulfonyl fluoride (PMSF), and 1 mM benzamidine on a nutating mixer for 30 minutes at 4°C. Cells were lysed using a Misonix S-4000 Sonicator with a ½” horn (Amplitude 20, 5 sec on, 10 sec off, 5 min total on-time). Cell lysate was clarified through ultracentrifugation at 160,000 X *g* for 45 minutes at 4°C in a Ti-70 rotor (Beckman-Coulter). Insoluble cell debris was removed and supernatant was filtered through a 0.2 μm polyethersulfone (PES) membrane bottle-top filter. Filtered lysate was loaded onto nickel nitrilotriacetic acid (Ni-NTA) Superflow resin (Qiagen) equilibrated with buffer 1 (50 mM Tris pH8, 500 mM NaCl, 10 mM imidazole). The column was washed with 500 ml buffer 1, followed by protein elution in 10 ml fractions of elution buffer 1 (50 mM Tris pH8, 500 mM NaCl, 250 mM imidazole). Elution fractions containing protein were dialyzed overnight at 4°C against 1.5 l dialysis buffer 1 (50 mM Tris pH8, 500 mM NaCl). Dialyzed protein was concentrated using a Vivaspin 20 3 kDa molecular weight cutoff (MWCO) concentrator (GE Healthcare) to 2 ml. The protein was then diluted to 100 mM NaCl immediately before loading onto a column packed with 10 ml High-S cation-exchange resin (Bio-Rad) attached to a Gilson Minipuls 3 peristaltic pump at 1.2 ml/min to reduce the likelihood of precipitation in low salt. The column was washed with 50 ml of low salt buffer (50 mM Tris pH8, 100 mM NaCl), followed by a 100 ml linear gradient from 100 mM to 1 M NaCl. Protein-containing fractions were pooled and dialyzed overnight at 4°C against 1.5 l dialysis buffer 1. Finally, the protein was concentrated to 0.7-1.2 mg/ml for crystallization experiments. Selenomethionine-labeled SeCsm2 (SeMet-SeCsm2) was purified using the same protocol.

### Recombinant SeCsm3 Protein Purification

Pellets were thawed on ice for 30 minutes before adding 1:1 volume of no salt buffer (50 mM Tris pH8, 20 mM imidazole). Resuspended cells were incubated with 50 μM lysozyme, 0.12% Brij 58, 1 mM PMSF, and 1 mM benzamidine on a nutating mixer for 30 minutes at 4°C. Additionally, 10 mM MgCl2, 2 mM CaCl2, and 16 μM bovine pancreas deoxyribonuclease I (Sigma Aldrich) was added and allowed to stir for 15 minutes at room temperature. Following this incubation, NaCl was added to a final concentration of 500 mM. Cells were further lysed using a Misonix S-4000 Sonicator with a ½” horn (Amplitude 20, 5 sec on, 10 sec off, 5 min total on-time). Insoluble cell debris was removed by ultracentrifugation at 160,000 X *g* for 45 minutes at 4°C in a Ti-70 rotor (Beckman-Coulter). To remove unbound nucleic acids, a 5% solution of polyethyleneimine (PEI) pH 7.9 was added dropwise to a final concentration of 0.2% into the supernatant while stirring at 4°C (Burgess, 1991). The solution was stirred for 5 minutes before removing the precipitated nucleic acids through centrifugation at 18,000 X *g* for 30 minutes in a FA-45-6-30 fixed-angle rotor (Eppendorf). The clarified lysate was filtered through a 0.2 μm PES membrane bottle-top filter. Filtered lysate was loaded onto a Ni-NTA Superflow resin column equilibrated with buffer 1. The column was washed with 200 ml buffer 1, followed by 400 ml buffer 2 (50 mM Tris pH8, 200 mM NaCl, 10 mM imidazole). Protein was eluted in 10 ml fractions with elution buffer 2 (50 mM Tris pH8, 200 mM NaCl, 250 mM imidazole). Protein-containing fractions were incubated with 1:10 molar concentration of tobacco etch virus (TEV) protease while dialyzing overnight at 4°C against 1.5 l dialysis buffer 2 (50mM Tris pH8, 100 mM NaCl, 10 mM imidazole). The dialysis bag was transferred to fresh 1.5 l dialysis buffer 2 and allowed to equilibrate again overnight at 4°C. Dialyzed, cleaved protein was subjected to another round of affinity chromatography to remove the cleaved His tag, uncleaved protein, and TEV protease. The Ni-NTA column flow through was loaded onto a High-S cation exchange resin column equilibrated with 50 mM Tris pH8, 100 mM NaCl at 0.4 ml/min. Nucleic acid-bound SeCsm3 binds to the resin, allowing the nucleic acid-free protein to be collected in the flow through. Cation exchange flow through was concentrated with a Vivaspin 20 10 kDa MWCO concentrator to 4-7 mg/ml to be used for crystallization experiments. SeMet-SeCsm3 was purified using the same protocol.

### Crystallization Conditions of SeCsm2 and SeCsm3

Crystallization trials of native and SeMet-derived SeCsm2 and SeCsm3 were conducted using a variety of commercial screens and utilizing both the hanging drop and sitting drop vapor diffusion methods. The best diffracting SeCsm2 and SeMet-SeCsm2 crystals were set up at 4°C in VDX hanging drop vapor diffusion plates (Hampton) and grown at 10°C in 100 mM Tris pH7, 8% ethanol at 1.2 mg/ml. Crystals form after 1-2 days of incubation. Crystals were cryo-protected by supplementing the crystallization condition with 20% ethylene glycol and 10% ethanol. SeMet-SeCsm3 crystals yielding the best diffraction data were set up at room temperature in Cryschem sitting drop vapor diffusion plates (Hampton) and grown at 14°C in 20% polyethylene glycol (PEG) 8000, 100 mM 2-(N-morpholino)ethanesulfonic acid (MES) pH6.5, 200 mM calcium acetate at 6.8 mg/ml. Crystals form after 24 hours. SeMet-SeCsm3 crystals were cryo-protected by supplementing the crystallization condition with 20% ethylene glycol. For heavy atom derivatization, SeMet-SeCsm3 crystals were soaked for 2 minutes in the cryo-protectant supplemented with 10 mM samarium (III) chloride before freezing.

### SeCsm2 Structure Determination

X-ray diffraction data were collected from native and SeMet SeCsm2 crystals at 100 K at the Life Sciences Collaborative Access Team (LS-CAT) beamlines at the Advanced Photon Source, Argonne National Laboratory, using a Rayonix MX300 CCD detector. All crystallographic data were processed with XDS (Kabsch, 2010) and Aimless (Evans & Murshudov, 2013). Positions of three Se atoms were obtained with the program Shake-and-Bake (Miller et al., 1993). A Single Anomalous Dispersion (SAD) electron density map was initially obtained using SHARP (delaFortelle & Bricogne, 1997). The map showed clearly the fold of the protein, but the data were limited to 3.2 Å resolution. An experimental electron density map including both the SeMet data and a higher resolution native data set was calculated with SHARP (delaFortelle & Bricogne, 1997) and was of better quality. The combined SeMet/native map allowed tracing of most of the molecule with the aid of Coot (Emsley & Cowtan, 2004) followed by iterative model building and refinement with BUSTER (Blanc et al., 2004), Phenix (Adams et al., 2010 ; Adams et al., 2011), and Refmac5 (Murshudov, Vagin, & Dodson, 1997). The final model spans most of the molecule, aside from one short disordered loop (residues 29-36) and two amino acids missing at the C-terminus, and includes 9 amino acids that form part of the His-tag used for purification. The stereochemistry of the model was validated with Molprobity (Chen et al., 2010) and Coot (Emsley & Cowtan, 2004). The final model has an R_work_/R_free_ of 24.47%/29.42% to 2.75 Å resolution with excellent stereochemistry. X-ray crystallographic statistics for the structure are summarized in **Supplementary Table 1**.

### SeCsm3 Structure Determination

X-ray diffraction data were collected from SeMet-SeCsm3 crystals at 100K at LS-CAT beamlines at the Advanced Photon Source, Argonne National Laboratory, using a Dectris Eiger 9M detector. Data from SeMet crystals soaked in 10 mM samarium (III) chloride were collected with the incident beam tuned to 11.5 keV (1.0781 Å) and 6.724 keV (1.8439 Å). Diffraction data were processed using XDS (Kabsch, 2010) and Aimless (Evans & Murshudov, 2013). An experimental electron density map was calculated using the automated pipeline CRANK-2 (Pannu et al., 2011) in CCP4 (Winn et al., 2011) using the 1.8439 Å wavelength data set. Initial model building was performed with CRANK-2 (Pannu et al., 2011) in CCP4 (Winn et al., 2011) followed by multiple iterations of automated model building using Buccaneer (Cowtan, 2006) and manual model editing using Coot (Emsley & Cowtan, 2004). The model was refined against the 1.0781 Å data set using Refmac5 (Murshudov et al., 1997). The final model contains two molecules in the asymmetric unit and spans most of the molecule, except for 3 disordered loops (A monomer 2-20, 32-65, 74-124, and 137-214, B monomer 0-20, 31-65, 75-124, 137-214), with an RMSD of 0.15 Å between the two molecules. The stereochemistry of the model was validated with MolProbity (Chen et al., 2010) and Coot (Emsley & Cowtan, 2004). The final model has an R_work_/R_free_ of 23.1%/26.7% to 2.40 Å resolution with excellent stereochemistry. X-ray crystallographic statistics for the structure are summarized in **Supplementary Table 2**.

### *S. epidermidis* RP62a Cas10-Csm Complex Purification

*S. epidermidis* RP62a harboring *pcrispr* with a Cas10 N-terminally His-tagged was generously provided by Dr. Asma Hatoum-Aslan at the University of Alabama (Hatoum-Aslan et al., 2013). Strains were grown in BBL brain heart infusion media (BD). The media was supplemented with neomycin (15 μg/ml) and chloramphenicol (10 μg/ml). Cultures (1 l culture per 2 l flask) were grown for 24 hours at 37°C. Cells were harvested, frozen at -80°C, and stored until purification. Cell pellets from 6 l of starting culture volume were thawed at 4°C for 30 minutes and resuspended in 30 ml of lysis buffer (30 mM MgCl_2_, 35 μg/ml lysostaphin [Ambi Products]) supplemented with PMSF and benzamidine (1 mM final concentration). Cells were incubated at 37°C for 1 hour, mixing every 20 minutes. Cell lysates were subsequently diluted 1:3 with resuspension buffer (200 mM NaH_2_PO_4_ pH7.4, 2 M NaCl, 80 mM imidazole, 0.4% Triton X-100, 2 mM PMSF, 2 mM benzamidine). Sonication was performed on the diluted lysates on ice (Amplitude 25, 5 sec on, 10 sec off, 10 minute total on-time [Misonix S-4000, ½” horn]). Sonicated lysate was spun at 18,000 X *g* for 45 minutes at 4°C in a FA-45-6-30 fixed-angle rotor. The insoluble cell debris was removed and spun again at 18,000 X *g* for 30 minutes. The remaining cell debris was discarded and the clarified lysate was filtered through a 0.2 μm PES membrane bottle-top filter. The filtered, clarified lysate was loaded onto a column packed with Ni-NTA Superflow resin (1 ml slurry per 1 l starting culture volume) equilibrated with equilibration buffer (50 mM NaH2PO4 pH7.4, 500 mM NaCl, 10 mM imidazole). The lysate was passed over the resin twice before washing with 600 ml wash buffer (50 mM NaH2PO4 pH7.4, 500 mM NaCl, 20 mM imidazole). Protein was eluted from the column with 25 ml elution buffer A (50 mM NaH2PO4 pH7.4, 500 mM NaCl, 250 mM imidazole) in 5 ml fractions. Protein-containing fractions were pooled and dialyzed against 1.5 l dialysis buffer A (20 mM Tris pH8, 200 mM NaCl) at 4°C overnight. Dialyzed complex was diluted to 100 mM NaCl immediately before loading onto a 10 ml column packed with Q Sepharose Fast Flow resin (GE Healthcare) equilibrated with buffer B (20 mM Tris pH8, 100 mM NaCl) running at 1.4 ml/min. The column was washed with 25 ml of buffer B, followed by a 100 ml linear gradient from 100 mM to 2 M NaCl. Fractions of interest were pooled and dialyzed against 1.5 l dialysis buffer B (20 mM Tris pH8, 500 mM NaCl) at 4°C overnight. Dialyzed complex was then concentrated to ~300 μl and loaded onto a Superdex 200 10/300 GL (GE Healthcare) equilibrated with dialysis buffer B. The non-aggregated complex peak was collected. Peak fractions are stored at 4°C for up to 1 month without any noticeable degradation as assayed by SDS-PAGE.

### Negative Stain Electron Microscopy Sample Preparation and Data Acquisition

Carbon-coated copper Gilder grids (300 mesh, Electron Microscopy Sciences [EMS]) were glow-discharged for 7 sec at 10 W in a Solarus 950 plasma cleaner (Gatan). Glow-discharged grids were placed carbon side down onto a 30 μl drop of 30 μg/ml purified complex for 5 minutes at room temperature. Grids were subsequently transferred twice to 30 μl drops of 20 mM Tris pH8, 500 mM NaCl for 1 minute per drop, with blotting occurring after the second transfer. The grids were then transferred twice to 30 μl drops of 2% uranyl acetate solution for 30 seconds per drop, followed by blotting after the second transfer and finally allowing the grid to air dry. Once dry, the grids were imaged on a JEOL 1400 transmission electron microscope (TEM) operating at 120 keV. 175 micrographs were collected using Leginon (Suloway et al., 2005) at a nominal magnification of 30,000x using an UltraScan4000 CCD camera with a pixel size of 3.71 Å at the specimen level and using a defocus range of -2 μm to -3 μm. Micrographs were processed using RELION-2.1 (Kimanius, Forsberg, Scheres, & Lindahl, 2016; Scheres, 2012). A small group of micrographs were selected for manually picking particles for initial 2D classification and the 2D classes were used for subsequent auto-picking of the entire dataset (~385,000 particles). Auto-picked particles were subjected to successive rounds of 2D classification to remove improperly picked particles yielding ~173,000 particles. The ~173,000 particles forming defined 2D classes were used in 3D classification. An input model was generated *ab initio* from a subset of these particles. The model best representing the 2D classes was chosen as the initial model for cryo-EM studies.

### Cryo-Electron Microscopy Sample Preparation and Data Acquisition

Cryo-EM grids (C-flat, 1.2/1.3, 400 mesh, Cu; [EMS]) were glow-discharged for 10 sec at 15 mA in an easiGlow glow discharger (Pelco). Grids were prepared under >95% humidity at 4°C using a Vitrobot Mark IV (FEI ThermoFisher). A glow-discharged cryo grid was placed in the Vitrobot sample chamber, onto which a 3 μl drop of protein (~0.2 mg/ml) was deposited. After waiting 5 seconds, the grid was blotted for 4 seconds with a blot force of +5 and immediately plunged into liquid ethane. The grid was transferred to a grid box for storage until data collection. Leginon (Suloway et al., 2005) was used for automated EM image collection. Micrographs were collected using a JEOL 3200FS TEM operating at 300 keV equipped with an in-column energy filter (Omega filter) and a K2 Summit direct electron detector (Gatan). A nominal magnification of 30,000x was used, for a pixel size of 1.24 Å at the specimen level and a defocus range of -2 μm to -3.5 μm. Movies were collected in counting mode with a total dose of ~37.5 e- per Å^2^, fractionated into 0.2 s frames for a total of 8 s, corresponding to 40 frames per movie. A total of 1,037 movies were collected in this manner.

### Cryo-EM Image Processing and Model Building

Cryo-EM movies were processed using RELION-3.0 (Nakane, Kimanius, Lindahl, & Scheres, 2018; Scheres, 2012; Zivanov et al., 2018). Motion correction was performed using the program MotionCor2 (Zheng et al., 2017) and dose-weighted according to the relevant radiation damage curves (Grant & Grigorieff, 2015) within RELION-3.0 (Zivanov et al., 2018) to allow for downstream post-processing. The Contrast Transfer Function (CTF) was estimated using CTFfind4 (Rohou & Grigorieff, 2015) on motion-corrected micrographs. 277,413 particles were auto-picked from templates generated from manual 2D classification. Auto-picked particles (277,413) were subjected to 3D classification, with the best class containing 107,620 particles. Additional RELION-3.0 post processing steps included per-particle CTF estimation and particle polishing to further improve the resolution. The particle polishing step showed a significant improvement on the final calculated volume, which had a 5.2 Å resolution according to the Fourier Shell Correlation (FSC) criterion (Rosenthal & Henderson, 2003). Initial inspection of the map showed that the quality was uneven, with the two extremes of the molecule at lower resolution than the central stem. The map was segmented into two pieces, the major stem (5 x SeCsm3 and 1 x SeCsm4), and the rest of the volume (putatively SeCas10), to use in multi-body refinement using RELION-3.0 (Nakane et al., 2018). Multi-body refinement did not improve the resolution of the stem, but did improve slightly the volume around the SeCas10 domain.

To position the SeCsm3 crystal structure in the cryo-EM map, a 6 dimensional search was done using the program Essens (Kleywegt & Jones, 1997), which identified 5 copies of SeCsm3 along the stem. The positions corresponded to the same locations identified visually. Searches using SeCsm2 as a template found no matches, indicating SeCsm2 was not part of the complex. Further visual inspection of the map showed the presence of an additional RRM motif at the base of the stem. Superposition of SeCsm3 on the RRM motif showed good agreement for the RRM region, but not for other regions, suggesting that the region corresponded to an RRM-containing protein distinct from SeCsm3. This was assigned as SeCsm4. The remaining density was assigned as SeCas10 based on the position and size. No density corresponding to SeCsm5 could be identified. Once all five SeCsm3 molecules had been identified, their position was refined using jigglefit (Brown et al., 2015) in Coot (Emsley & Cowtan, 2004). The models showed excellent fit in most areas, but some loops and secondary structure elements needed adjustment to fit the density. To improve the placing of the crystal structures, the monomers were fit into the map using the MDFF routines (Trabuco et al., 2008) that are part of NAMD (Phillips et al., 2005). No further refinement of the models was done. Unless noted, the figures were made using the crystal structures fit into the map.

### Model superpositions, comparisons, and figures

Comparisons of the different models were all done using Coot (Emsley & Cowtan, 2004) and Chimera (Pettersen et al., 2004). When superposing structures from different organisms, the superposition was based on secondary structure matching (SSM) as implemented in Coot (Emsley & Cowtan, 2004) and Chimera (Pettersen et al., 2004).

Figures were drawn with Pymol (Pymol) and Chimera (Pettersen et al., 2004). Conservation data were generated using the ConSurf server (Ashkenazy et al., 2010; Landau et al., 2005) and visualized in Pymol. Electrostatic potential calculations were done with APBS (Baker, Sept, Joseph, Holst, & McCammon, 2001).

## Acknowledgements

We thank members of the Mondragón laboratory for discussions and assistance. We thank Jonathan Remis for help with cryo-EM data collection. We acknowledge help and discussions with Erik Sontheimer in the early stages of the project. Research was supported by the NIH (A.M., grant R35-GM118108). We acknowledge the help from the Northwestern University Structural Biology Facility, Keck Biophysics Facility, and High Throughput Analysis Laboratory. Support from the R.H. Lurie Comprehensive Cancer Center of Northwestern University to the Structural Biology Facility is acknowledged. LS-CAT/Sector 21 at the Advanced Photon Source, Argonne National Laboratory was supported by the Michigan Economic Development Corporation and the Michigan Technology Tri-Corridor.

---

**Supplementary Table 1.** **X-ray crystallographic data statistics for SeCsm2 crystals**

|  | **Native** | **SeMet** |
| --- | --- | --- |
| **Data Collection** |  |  |
| Detector type/source | RayonixCCD/APS | RayonixCCD/APS |
| Wavelength (Å) | 0.97856 | 0.97872 |
| Resolution range* (Å) | 77.78-2.75 (2.88-2.75) | 39.40-3.2 (3.42-3.20) |
| Space group | P4<sub>1</sub>2<sub>1</sub>2 | P4<sub>1</sub>2<sub>1</sub>2 |
| a, b, c (Å) | 77.78, 77.78, 69.47 | 78.8, 78.8, 68.27 |
| Measured reflections* | 39,163 (3,861) | 34,368 (6,065) |
| Unique reflections* | 5,931 (765) | 3,852 (665) |
| Completeness* (%) | 99.9 (99.8) | 99.2 (96.3) |
| Mean [I/σ(I)]* | 22.7 (1.8) | 13.8 (1.4) |
| Multiplicity* | 6.6 (5.0) | 8.9 (9.1) |
| R<sub>measure</sub>* | 0.052 (0.956) | 0.163 (1.94) |
| CC(1/2)* | 0.999 (0.645) | 0.998 (0.527) |
| MFID<sup>^</sup> | - | 0.244 (0.375) |
| **Phasing** |  |  |
| SeMet Sites | - | 3 |
| Phasing Power | - |  |
| Dispersive (centric/accentric) | - | 0.717/0.802 |
| Anomalous (acentric) | - | 1.301 |
| FOM (centric/accentric) | - | 0.224/0.157 |
| **Refinement** |  |  |
| Number of reflections working/test | 5,635/256 | - |
| R (working set; %) | 24.47 | - |
| R<sub>free</sub> (%) | 29.42 | - |
| **Structure Quality** |  |  |
| Protein atoms | 1,071 | - |
| RMS deviations in bond lengths (Å) | 0.0144 | - |
| RMS deviations in bond angles (°) | 1.69 | - |
| Ramachandran plot - most favored regions (%) | 98.36 | - |

\* All numbers in parenthesis are for highest resolution shell
<sup>^</sup> Mean fractional isomorphous difference against Native

**Supplementary Table 2.** **X-ray crystallographic data statistics for SeCsm3 crystals**

|  | Remote | Edge |
| --- | --- | --- |
| **Data Collection** |  |  |
| Detector type/source | Dectris Eiger 9M/APS | Dectris Eiger 9M/APS |
| Wavelength (Å) | 1.07810 | 1.8439 |
| Resolution range* (Å) | 46.57 – 2.40 (2.49 – 2.40) | 51.9 – 2.33 (2.46 – 2.33) |
| Space group | C2 |  |
| a, b, c (Å) | 104.6, 50.98, 83.02 | 104.6, 50.75, 83.31 |
| $\beta$ (°) | 97.05 | 97.13 |
| Measured reflections* | 114,637 (10,902) | 311,330 (11,697) |
| Unique reflections* | 16,891 (1,698) | 16,657 (1,328) |
| Completeness* (%) | 98.2 (96.1) | 88.9 (49.3) |
| Mean $[I/\sigma(I)]^*$ | 10.3 (2.0) | 10.2 (0.9) |
| Multiplicity* | 6.8 (6.4) | 18.7 (8.8) |
| $R_{\text{measure}}^*$ | 0.126 (0.951) | 0.268 (2.531) |
| CC(1/2)* | 0.996 (0.741) | 0.996 (0.501) |
| **Phasing** |  |  |
| Sm Sites |  | 8 |
| FOM |  | 0.194 |
| **Refinement** |  |  |
| Number of reflections working/test | 16,078/795 | - |
| $R$ (working set; %) | 23.05 (30.5) | - |
| $R_{\text{free}}$ (%) | 26.69 (39.9) | - |
| **Structure Quality** |  |  |
| Protein atoms | 2,932 | - |
| Samarium ions | 13 | - |
| Waters | 61 | - |
| Other | 10 | - |
| RMS deviations in bond lengths (Å) | 0.009 | - |
| RMS deviations in bond angles (°) | 1.292 | - |
| Ramachandran plot - most favored regions (%) | 96.3 | - |

\* All numbers in parenthesis are for highest resolution shell

**Supplementary Table 3.** **SeCsm3 Residues that have been characterized biochemically**.

| **Function** | **Residue** | **Reference** |
| --- | --- | --- |
| crRNA Binding & Maturation | K52 | (Walker et al., 2017) |
| " | K54 | " |
| " | R56 | " |
| " | D100 | (Hatoum-Aslan et al., 2013) |
|  | E120 |  |
|  | E124 |  |
|  | E140 | (Walker et al., 2017) |
| Protein stability | G182 | (Hatoum-Aslan et al., 2014) |
| " | G183 | " |
| " | G185 | " |
| Csm3-Csm3 Interface | K4 | (Walker et al., 2017) |
| " | D179 | " |
| Target RNA Cleavage | H18 | (Hatoum-Aslan et al., 2013) |
| " | D32 | (Samai et al., 2015) |

**Supplementary Table 4.**
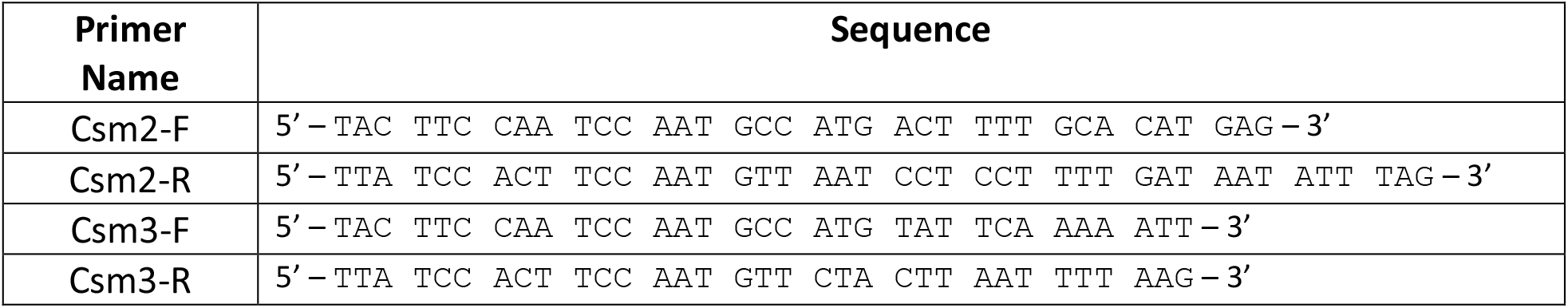
**Primers used for cloning SeCsm2 and SeCsm3**

**Figure 2 - figure supplement 1.**
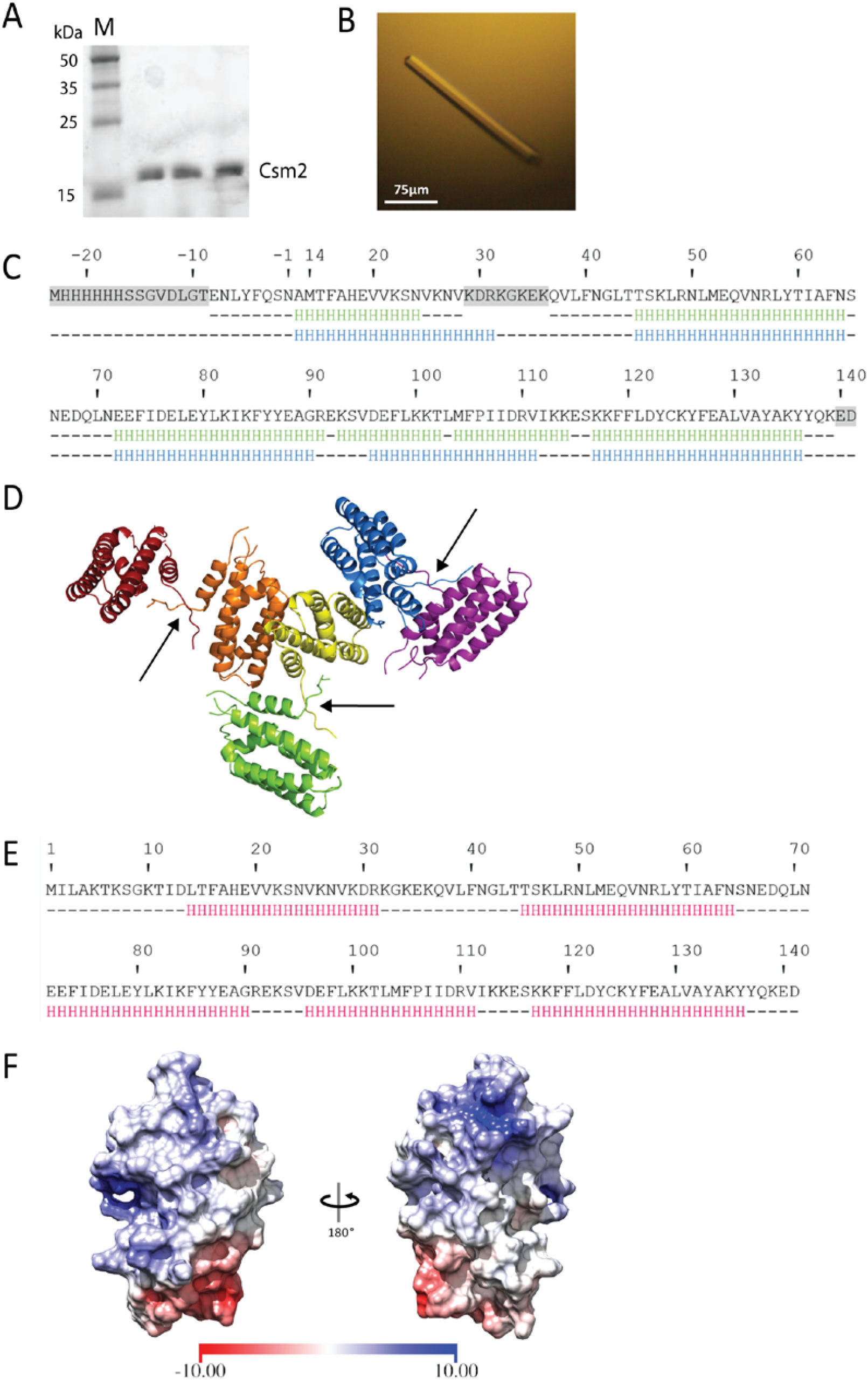
Purification and Structure Determination of SeCsm2. (A) Coomassie blue stained 10% SDS-PAGE of purified SeCsm2 after the final cation exchange column. The three lanes correspond to three different fractions. M, molecular weight markers. (B) Photograph of a representative crystal of SeCsm2. SeCsm2 forms square-shaped rod-like crystals with dimensions ~200 μm x ~20 μm x ~20 μm. (**C**) Amino acid sequence of the SeCsm2 construct used in the crystallization experiments. Disordered regions in the crystal structure are highlighted in gray. The secondary structure observed in the crystal (green) and the predicted secondary structure (blue) are shown below the sequence, where H denotes an α-helical residue. The greatest difference between the actual and predicted secondary structures lies at α_4_-α_5_ in the crystal sequence (α_4_ in prediction). (**D**) Schematic diagram of the crystal packing of SeCsm2 showing critical lattice contacts formed between the affinity tag regions of adjacent molecules, explaining why the affinity tag was necessary for crystallization. (**E**) SeCsm2 amino acid sequence using the revised gene annotation and with an updated secondary structure prediction (red). The N-terminal region not part of the crystal structure is not predicted to form a secondary structure element. (**F**) SeCsm2 accessible surface colored by electrostatic potential calculated with APBS (Baker et al., 2001). Two small positive regions (blue) are found on opposite sides of the molecule. One negative region (red) is shown at the opposite end from the positive regions. The bar at the bottom maps the electrostatic potential values from -10 kBT/ec [red] to 10 kBT/ec [blue].

**Figure 3 – figure supplement 1.**
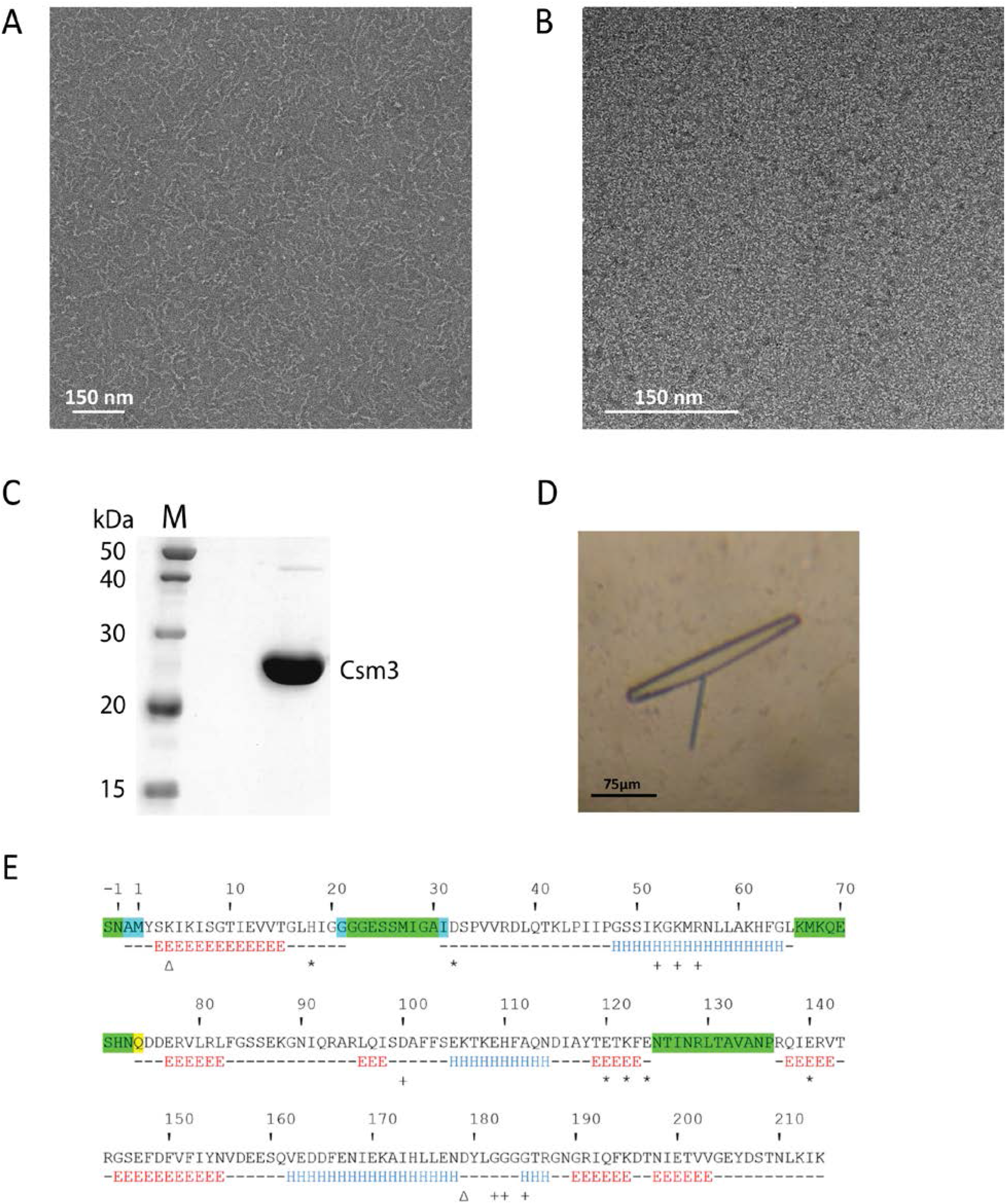
Purification and Sequence of SeCsm3. (**A**) Negative stain image of nucleic acid-bound SeCsm3. SeCsm3 forms a characteristic spiral shape when bound to nucleic acids. Negative stain image taken at 40,000x magnification. (**B**) Negative stain image of nucleic acid-free SeCsm3. Nucleic acid-free SeCsm3 does not show any aggregation or spiral formation. The lack of aggregation is required for crystallographic studies. Negative stain image taken at 100,000x magnification. (**C**) Coomassie blue stained 12% SDS-PAGE of SeCsm3 after the final cation exchange column. M, molecular weight markers. (**D**) Photograph of a representative crystal of SeCsm3. SeCsm3 forms thin, overlapped plates with dimensions ~175 μm x ~15 μm ~10 μm. Data were collected only on non-overlapping regions of the crystals. (**E**) SeCsm3 amino acid sequence of the construct used in crystallography studies. Disordered regions in both chain A and B are shown in green, with additional residues missing from chain A shown in blue and chain B in yellow. Chain B was used in all data analyses and diagrams. The secondary structure within the crystal is shown below the sequence. H, α-helical residue (blue); E, residue in β-strand (red). Biochemically studied residues are labeled based on relation to crRNA binding (+), target RNA binding and cleavage (*), and interface interactions (Δ).

**Figure 3 – figure supplement 2.**
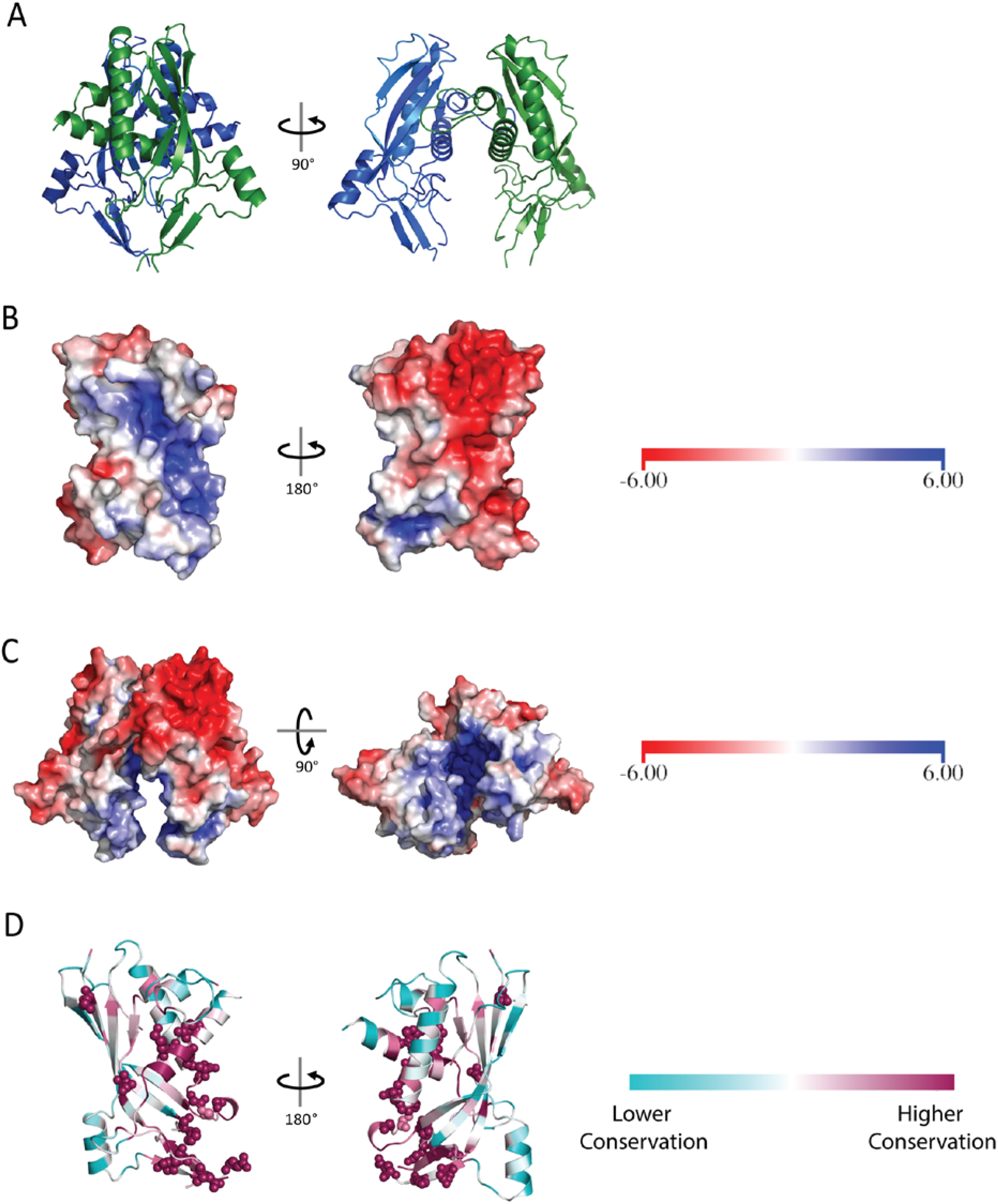
Crystallographic and Amino Acid Conservation Analysis of SeCsm3. (**A**) Schematic diagrams of two orthogonal views of the SeCsm3 dimer. The SeCsm3 crystallographic dimer shows contacts between monomers involving a loop and a helix. (**B**) SeCsm3 solvent accessible surface colored by electrostatic potential. The surface shows a large, highly positively charged region (blue) on one side of the molecule, with the opposite side of the molecule more negatively charged (red). (**C**) Accessible surface of the SeCsm3 crystallographic dimer colored by electrostatic potential. The positively charged region (blue) is present at the interface of the dimer, partially shielding this region from the solvent. In **B** and **C**, the bars on the right hand side map the electrostatic potential values from -6 kBT/ec [red] to 6 kBT/ec [blue]. (**D**) Schematic diagram of SeCsm3 colored by amino acid sequence conservation. Analysis of residue conservation shows a clustering of highly conserved residues corresponding to the positively charged region in **B**. Biochemically studied residues (Hatoum-Aslan et al., 2014; Hatoum-Aslan et al., 2013; Samai et al., 2015; Walker et al., 2017) are shown as spheres, with all being highly conserved. These residues have been implicated in functions such as crRNA binding, target RNA binding and cleavage, and intermolecular interactions. Amino acid conservation was calculated using the ConSurf server (Ashkenazy et al., 2010; Landau et al., 2005). Highly conserved residues are shown in maroon with less conserved residues in cyan.

**Figure 5 – figure supplement 1.**
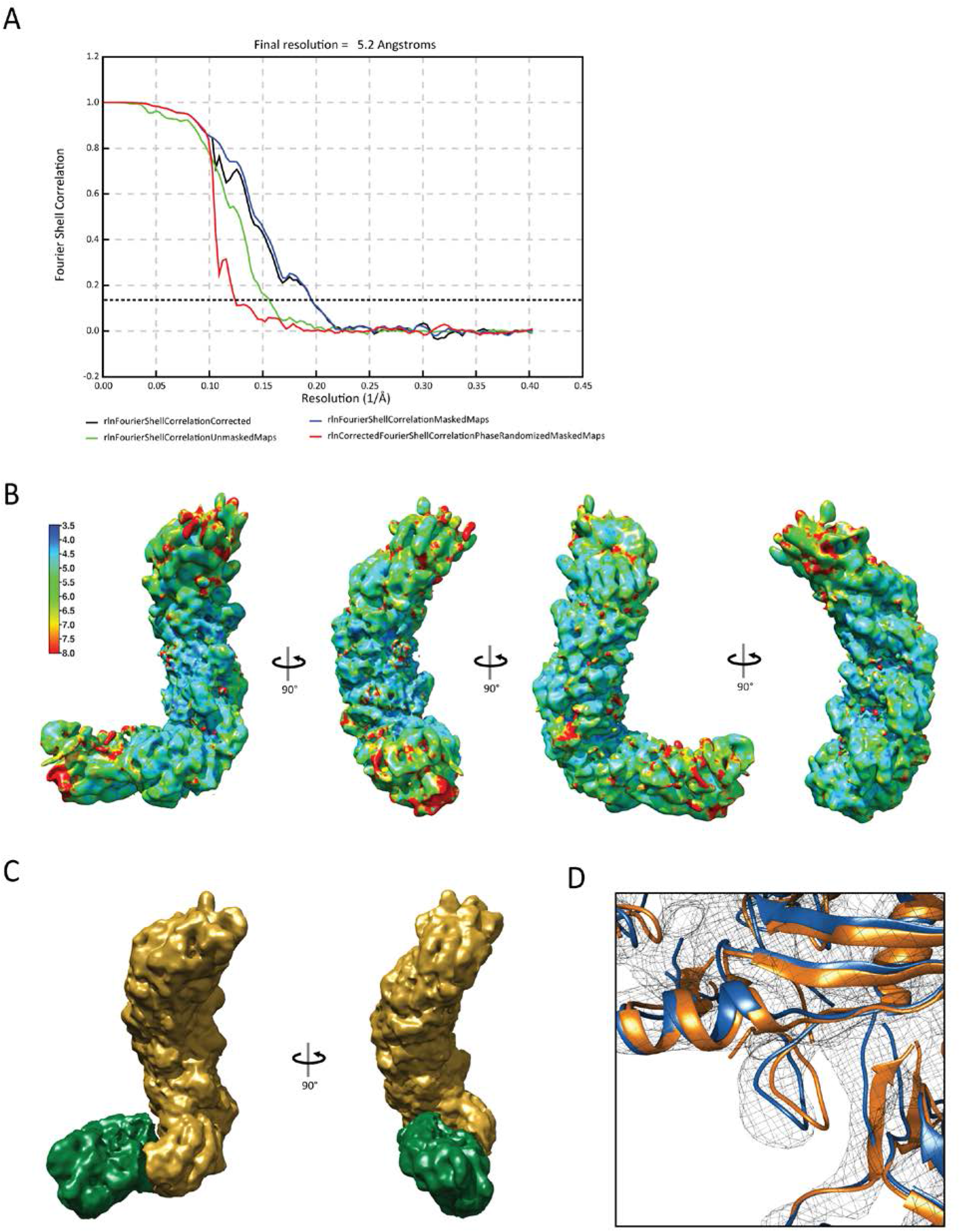
Cryo-EM Structural Analysis of the *S. epidermidis* Effector Subcomplex. (**A**) Fourier Shell Correlation (FSC) curve for the final cryo-EM volume. The dashed line corresponds to a correlation of 0.143. The estimated resolution is 5.2 Å using the 0.143 gold-standard FSC criterion implemented in RELION (Scheres, 2012, 2016; Zivanov et al., 2018). (**B**) Local resolution estimation (Kucukelbir et al., 2014) of the SeCas10-Csm subcomplex shows that the stem has the highest resolution while both ends of the subcomplex show a lower resolution, probably due to conformational heterogeneity. The bar on the left maps resolution estimates to color in the map. (**C**) Diagram of the segmented volumes of the stem (yellow) and base (green) generated for use in multi-body refinement (Nakane et al., 2018). (**D**) Flexible fitting of SeCsm3 into the cryo-EM density with MDFF (Trabuco et al., 2008) (blue) shows minimal rearrangement of the secondary structure elements, but larger conformational rearrangements of the exposed loops within SeCsm3. The Essens (Kleywegt & Jones, 1997) rigidly docked SeCsm3 subunits are shown in orange.

**Figure 6 – figure supplement 1.**
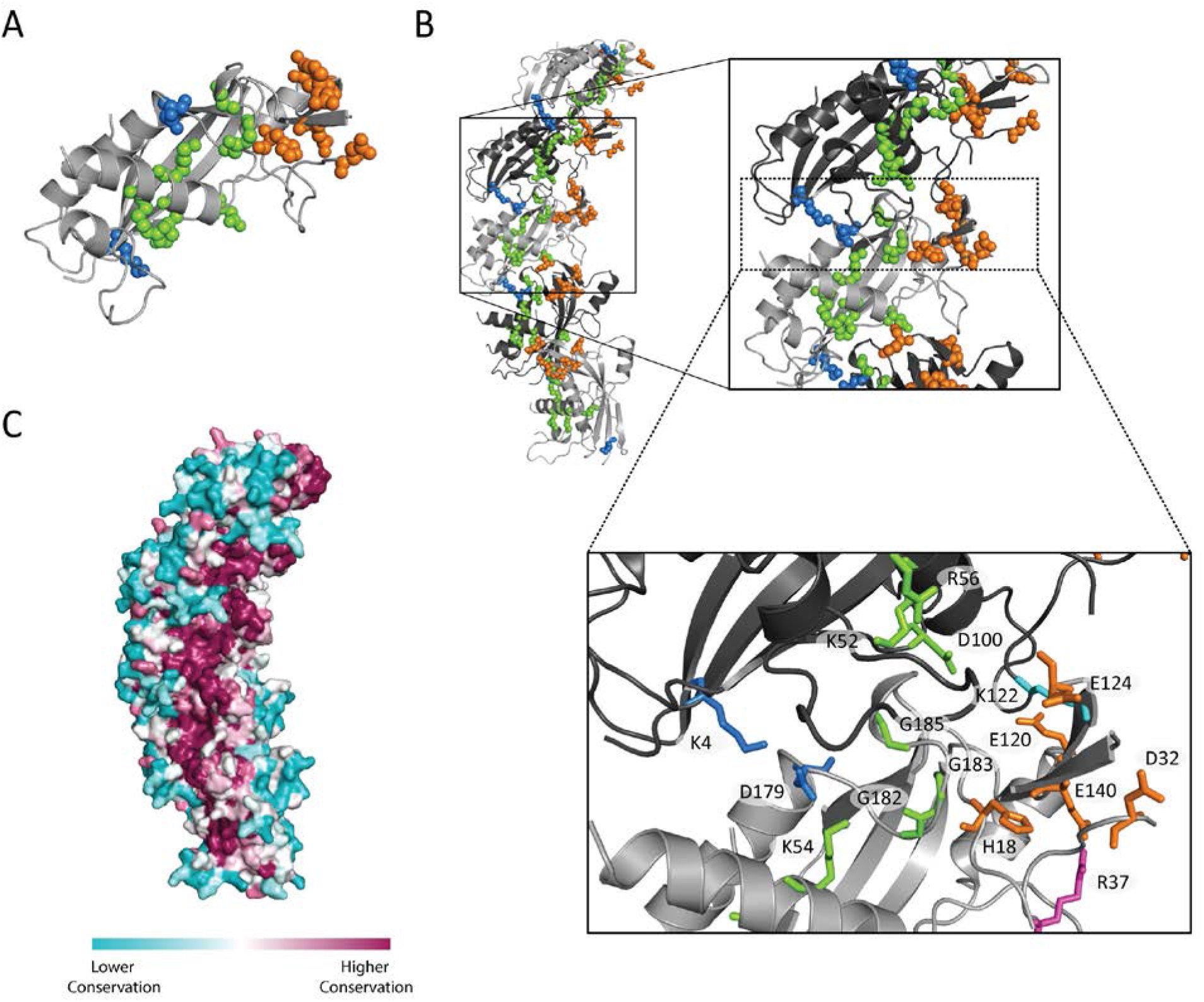
Location in SeCsm3 in the Subcomplex of Biochemically Analyzed Amino Acids. (**A**) Amino acid residues in SeCsm3 previously studied biochemically (Hatoum-Aslan et al., 2014; Hatoum-Aslan et al., 2013; Samai et al., 2015; Walker et al., 2017) are shown as spheres in the schematic diagram of a SeCsm3 monomer. Residues are colored based on their role in crRNA binding (green), target RNA binding and cleavage (orange), or SeCsm3-SeCsm3 interface interactions (blue) (**Supplementary Table 3**). (**B**) Schematic diagram of the SeCas10-Csm subcomplex showing adjacent SeCsm3 subunits with a zoomed region of two adjacent SeCsm3 subunits, which shows spatial segregation of the residues with different functional roles, colored as in **A**. Additionally, two residues, R37 (magenta) and K122 (cyan), are shown as they may help stabilize the interface interacting with previously identified residues. The diagram shows that many of the highlighted residues interact across two adjacent subunits, while others lie within the positively charged channel in the stem. (**C**) Surface representation of the SeCsm3 stem colored by sequence conservation calculated using the ConSurf server (Ashkenazy et al., 2010; Landau et al., 2005). Highly conserved residues (magenta) correlate well to the location of the positive channel, with less conserved residues (cyan) on the exterior of the stem.

**Figure 6 – figure supplement 2.**
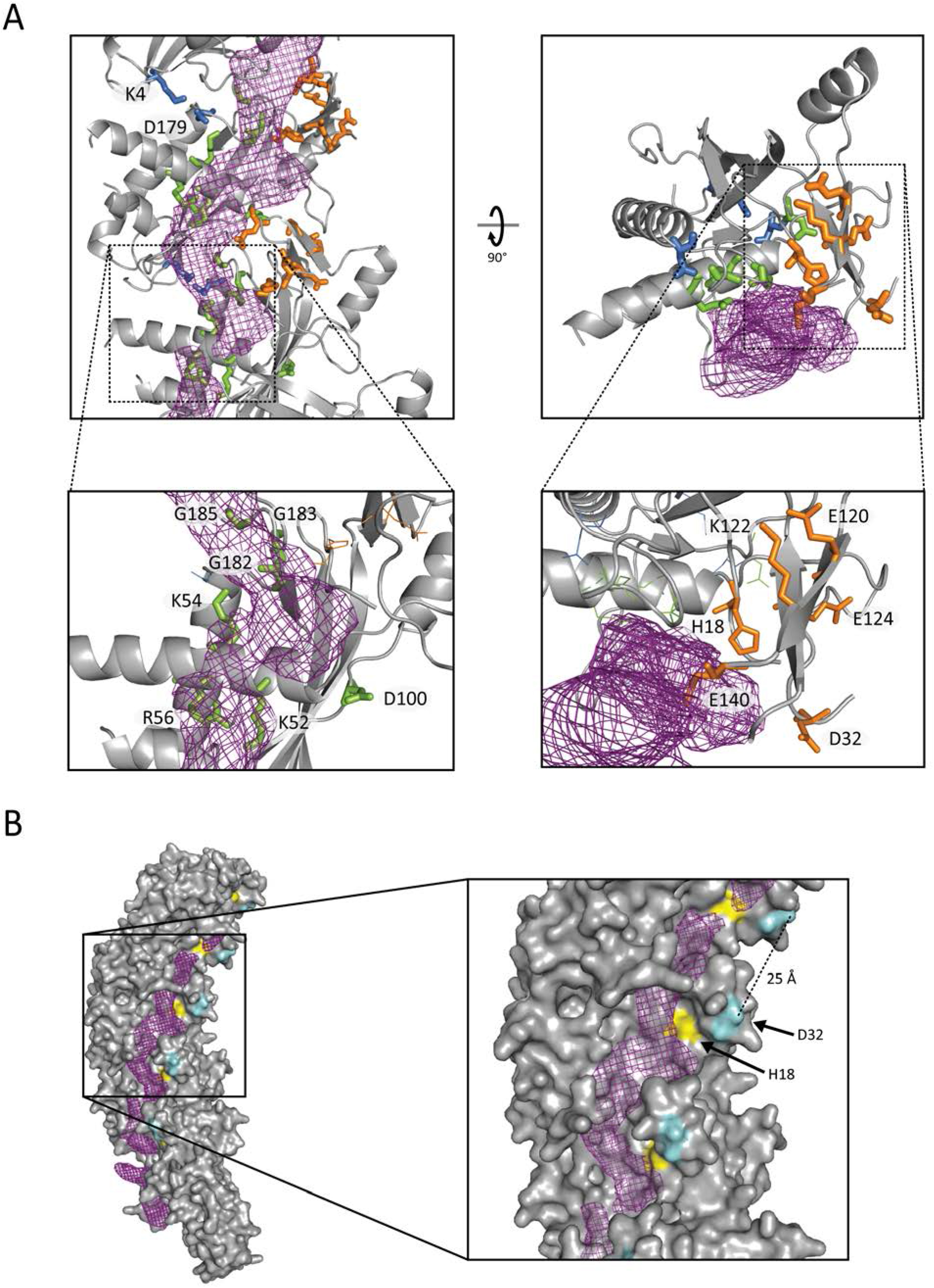
Location of crRNA and its Relation to Biochemically Analyzed Amino Acids in the Subcomplex. (**A**) Schematic diagram of orthogonal views of the SeCas10-Csm subcomplex with the density assigned to the crRNA overlaid. The location of the density shows that the residues involved in crRNA binding (green) are found near the crRNA density. The residues associated with target RNA binding and cleavage (orange) are distant from the crRNA density. (**B**) Diagram of the accessible surface of the SeCas10-Csm subcomplex with the crRNA density overlaid. The diagram shows that the distance between putative active site residues on adjacent SeCsm3 subunits is consistent with an extended target RNA conformation. Aspartate 32 (D32, blue) marks the location of the target RNA cleavage active site. A conserved histidine (yellow) has been implicated in target RNA cleavage and lies close to the putative active site centered on residue D32.

**Figure 6 – figure supplement 3.**
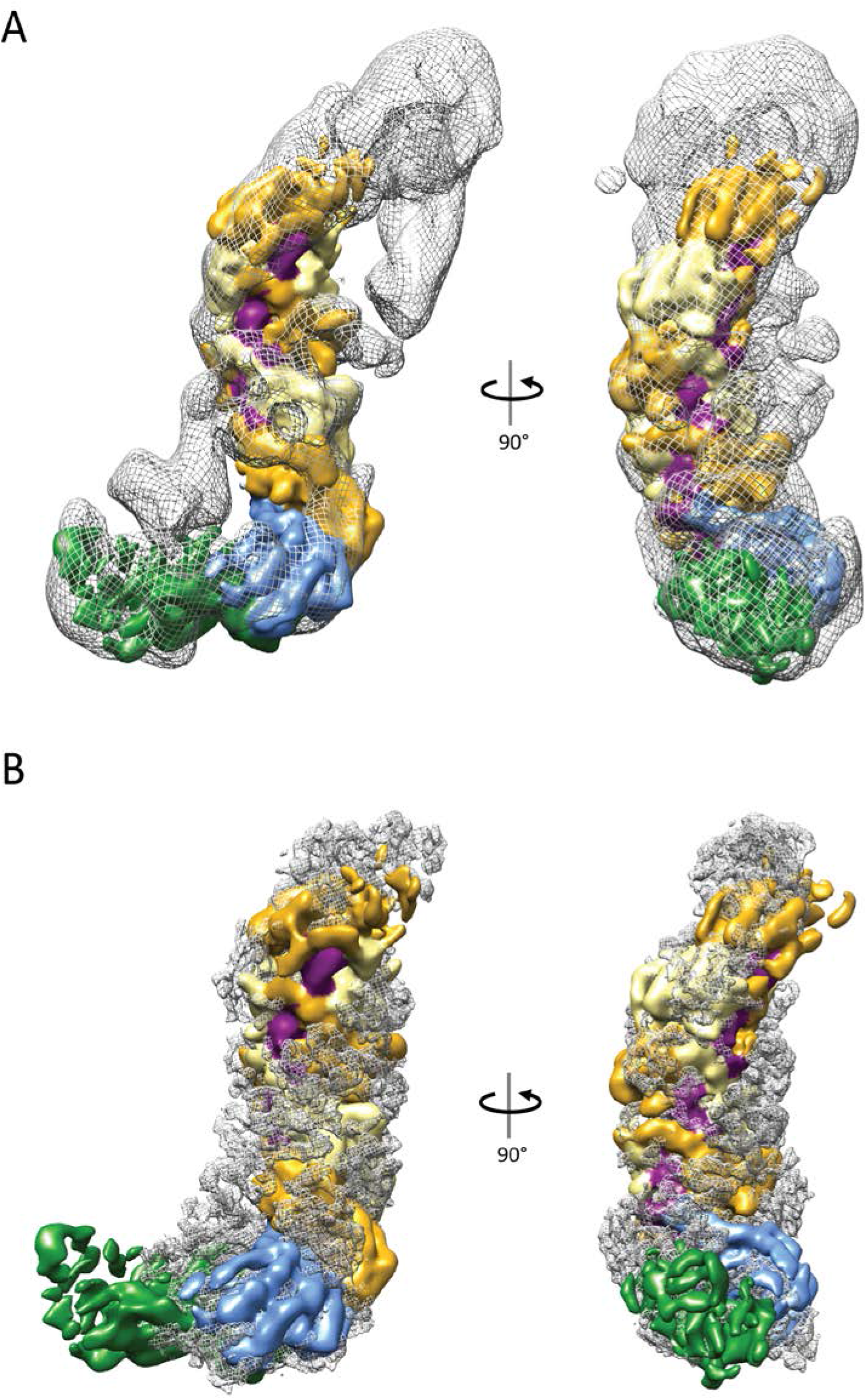
Structural Comparison of the SeCas10-Csm Subcomplex with the *T. thermophilus* Csm (Type III-A) and Cmr (Type III-B) Complexes. (**A**) Diagram showing orthogonal views of the alignment of the *T. thermophilus* Cas10-Csm complex reconstruction (17 Å resolution; EMD-6122) (R. H. Staals et al., 2014) (mesh) with the SeCas10-Csm subcomplex reconstruction (solid surface colored as in **Figure 6A**). The alignment shows the strong similarity between the two complexes. In the TtCas10-Csm effector complex, the density at the top was assigned to Csm5 and three Csm2 were assigned to the minor helical filament adjacent to the Csm3 major filament. (**B**) Diagram showing orthogonal views of the alignment of the *T. thermophilus* Cmr complex reconstruction (4.1 Å resolution; EMD-2898) (Taylor et al., 2015) (mesh) with the SeCas10-Csm subcomplex reconstruction (solid surface colored as in **Figure 6A**). The alignment shows that the TtCas10-Cmr complex is a more cylindrical complex without a protrusion at the base. Differences in subunit structures and stoichiometry also contribute to the overall structural variations between the two complexes.

